# The E3 ubiquitin ligase HECTD1 contributes to cell cycle progression through an effect on mitosis

**DOI:** 10.1101/2021.12.17.473173

**Authors:** Natalie Vaughan, Nico Scholz, Catherine Lindon, Julien D. F. Licchesi

## Abstract

Mechanistic studies of how protein ubiquitylation regulates the cell cycle, in particular during mitosis, has provided unique insights which have contributed to the emergence of the ‘Ubiquitin code’. In contrast to RING E3 ubiquitin ligases such as the APC/c ligase complex, the contribution of other E3 ligase families during cell cycle progression remains less well understood. Similarly, the contribution of ubiquitin chain types beyond homotypic K48 chains in S-phase or branched K11/K48 chains assembled by APC/c during mitosis, also remains to be fully determined. Our recent findings that HECTD1 ubiquitin ligase activity assembles branched K29/K48 ubiquitin linkages prompted us to evaluate its function during the cell cycle. We used transient knockdown and genetic knockout to show that HECTD1 depletion in HEK293T and HeLa cells decreases cell proliferation and we established that this is mediated through loss of its ubiquitin ligase activity. Interestingly, we found that HECTD1 depletion increases the proportion of cells with aligned chromosomes (Prometa/Metaphase). We confirmed this molecularly using phospho-Histone H3 (Ser28) as a marker of mitosis. Time-lapse microscopy of NEBD to anaphase onset established that HECTD1-depleted cells take on average longer to go through mitosis. To explore the mechanisms involved, we used proteomics to explore the endogenous HECTD1 interactome in mitosis and validated the Mitosis Checkpoint Complex protein BUB3 as a novel HECTD1 interactor. In line with this, we found that HECTD1 depletion reduces the activity of the Spindle Assembly Checkpoint. Overall, our data suggests a novel role for HECTD1 ubiquitin ligase activity in mitosis.

## 1. Introduction

The Ubiquitin-Proteasome System (UPS) plays important roles during the cell cycle through targeted degradation of key checkpoint proteins. In mitosis, the turnover of cyclins is required in order to commit cells to anaphase and complete cell division. This is achieved through the highly conserved post-translational modifier ubiquitin [1]. Ubiquitin is covalently attached, mostly onto exposed lysines residue of proteins, through a multi-step mechanism which require the activity of an E1-activating enzyme [2–4], E2-conjugating enzymes and E3 ubiquitin ligases [5, 6]. E3 ubiquitin ligases such as Really Interesting New Gene (RING) function as scaffold, bringing the ubiquitin loaded-E2 into proximity to the substrate to enable ubiquitin transfer. The E2-conjugating enzyme UBE2S for example cooperates with the Anaphase Promoting Complex/cyclosome (APC/c) to extend K11-linked ubiquitin chains onto mitotic substrates, triggering mitotic exit and signalling for the completion of cell division [7–11].

Ubiquitin can be attached onto protein targets as a single moiety or as a ubiquitin chain. It is assembled into chains through an isopeptide bond between any of its seven lysines (K6, K11, K27, K29, K33, K48 and K63) or the N-terminus (i.e., linear ubiquitin) of an acceptor ubiquitin and the C-terminus of a donor ubiquitin, with all these chain types being found in eukaryotes [12, 13]. Therefore, as many as eight linkage types can be used to assemble homotypic, heterotypic, mixed and even branched ubiquitin chains onto protein targets [14]. These ubiquitin linkages have different structural and biochemical properties and indeed cellular functions. The cell cycle is perhaps the best example of how these different signals can work in concert for the temporal regulation of molecular mechanisms [15, 16]. K48 and K11-linked chains have been shown to function as efficient degradation signals by the UPS, allowing for the timely turnover of cell cycle proteins. For example, K48-linked ubiquitin chains mediate protein degradation of Cyclin Dependent Kinase inhibitor p21, the kinase PLK4 and the phosphatase CDC25 during S-phase. In contrast, the APC/c ubiquitin ligase complex polyubiquitinates cyclin B1, securin as well as aurora kinase A and B with K11-linked chains and thus targets these mitotic proteins for subsequent degradation by the UPS [17–23].

Although most of our understanding of ubiquitin biology relates to homotypic chains, atypical and branched ubiquitin chains have been recently emerged as important signals for cell cycle regulation. For example, branched K11/K48 chains assembled by APC/c have emerged as improved signals for the efficient degradation of mitotic cyclins [8, 11, 24, 25]. In addition to RING and Skp1-Cul1-F-box protein (SCF) E3s, HECT E3s have also been implicated in cell cycle regulation [26, 27]. In contrast to RING E3s, the Homologous to the E6AP C-terminus (HECT) family of E3 ligases are characterised by intrinsic enzyme activity contributed by a catalytic cysteine residue located in the C-lobe, and which accepts ubiquitin from the loaded-E2 through trans-thioesterification prior to transfer onto lysine residue(s) of protein substrates [28–30]. SMURF2 belongs to the NEDD4 subfamily of HECT E3 ubiquitin ligases and was first identified as a regulator of BMP/TGF-β signalling prior to being implicated in chromatin organisation and mitotic regulation. SMURF2 deletion for example impairs the Spindle Assembly Checkpoint (SAC) resulting in defective chromosome alignment/segregation and premature anaphase onset [27]. Further mechanistic studies established SMURF2 uses its ligase activity to assemble non-degradative K63-linked chains onto the Mitosis Checkpoint Complex protein MAD2, thereby protecting it from K48-linked polyubiquitylation and subsequent degradation by the 26S proteasome. SMURF2 has also been implicated at the G2/M transition where its ligase activity stabilises NEDD9 which is required for Aurora A activation and mitotic entry [31]. These studies highlight how the same E3 ubiquitin ligase can act at different stages during the cell cycle and further argues that the ‘ubiquitin code’ in mitosis is more complex than originally thought, with various degradative, non-degradative, typical, and atypical ubiquitin signals involved. In further support of this, the HECT E3 ligase TRIP12 which participates in the DNA damage response pathway also employs a mechanism independent of its ubiquitin ligase activity to regulate mitotic entry by controlling DNA replication, mitotic progression, and chromosome stability [32, 33].

TRIP12 as well as its yeast ancestor UFD4 can assemble homotypic K29-linked ubiquitin onto protein targets but also onto pre-assembled ubiquitin chains [34, 35]. Although HECTD1, the closest homologue to TRIP12, can also use K29 to build chains, it preferentially assembles branched K29/K48-linked ubiquitin chains, *in-vitro* at least [36]. Interestingly, TRIP12, UFD4 and HECTD1 are involved in the DNA damage response and chromatin regulation, with HECTD1 recently implicated in histone ubiquitylation during Base Excision Repair (BER) [37, 38]. Yet, whether this effect is mediated through its activity towards K29/K48 branched ubiquitin chains remains to be shown. In addition, multiple functions for HECTD1 have been proposed including during embryonic development, the regulation of cell migration as well as in various signal transduction pathways such as Wnt, Retinoic acid signalling and NFK κ-B [39–46]. Different types of ubiquitin chains have been ascribed to HECTD1, with K48-linked ubiquitin chains implicated in the regulation of the focal adhesion protein ACF7 and PIPKIγ90, while K63 has been suggested for its role on HSP90 and the Wnt antagonist Adenomatous polyposis coli (APC) [41, 47, 48]. Further, HECTD1 has also been implicated in regulating Estrogen Receptor (ER)-mediated transcription. In this mechanism, condensin I and condensin II are recruited to the ER-α enhancers, and in turn recruit HECTD1, which lead to the ubiquitin-dependent degradation of the corepressor RIP40 and transcriptional activation of ER-α gene targets [49]. HECTD1 therefore appears to assemble both proteasomal and non-proteasomal ubiquitin signals, and it will be important to delineate the contribution of its K29/K48 ligase activity on its cellular functions.

In this study we explored HECTD1 function in the context of cell proliferation and found that loss of HECTD1 ubiquitin ligase activity reduces cell proliferation through an effect on mitosis. Together, our data support recent evidence implicating K29-linked ubiquitylation in mitotic regulation and provide novel insights into the ‘ubiquitin code’ [50].

## 2. Materials and Methods

### 2.1. Mammalian cell culture

Human embryonic kidney 293T (HEK293T) cells, HEK293T HECTD1 knock out cells (Dr Bienz, MRC-LMB Cambridge, UK), HEK293ET and HeLa cells were cultured in Dulbecco’s modified minimum essential medium (DMEM) with GlutaMAX supplement, supplemented with 10% (v/v) Fetal Bovine Serum and 1% (v/v) 10,000 units Penicillin-10mg/ml Streptomycin, at 37°C in a humidified incubator with 5% CO_2_ [51]. Cells were passaged by incubating with sterile 0.05% EDTA-PBS for 5 min at 37°C, followed by pelleting the cells at 1,000 rpm for 3 min. Cells were then resuspended in supplemented DMEM (with GlutaMAX) and seeded (1/10) in a Nunclon™ Delta surface-treated (Nunc™) 10 cm dish. HECTD1 knock out cells were cultured from passage 4 to passage 35. Both HEK293T HECTD1 KO1 and KO2 have been generated using the same gRNA and these are confirmed individual clones (and not pools) [51]. Glioblastoma cell lines U87 and U251 were purchased from ATCC and cultured in Eagle’s Minimum Essential Media (EMEM) (Sigma-Aldrich, M2279), supplemented with 10% (v/v) Fetal Bovine Serum (Heat-inactivated FBS), 2 mM L-glutamine, MEM 1% (v/v) non-essential amino acids, 1 mM sodium pyruvate, and 1%(v/v) 10,000 units Penicillin-10 mg/ml Streptomycin, at 37°C in a humidified incubator with 5% CO_2_. U87 and U251 cell lines were passaged as described above. All cell lines were tested for Mycoplasma contamination, using the MycoAlert™ Mycoplasma Detection Kit (Lonza, Switzerland), as per manufacturer’s instructions.

### 2.2. Cell synchronisation

To synchronise the cells with RO3306, cells were treated with 9 μM RO3306 for 20 hrs. To release cells into mitosis, cells synchronised with RO3306 were washed three times in PBS, and released into fresh media.

For double thymidine block, cells were treated with 2 mM thymidine (Sigma-Aldrich, T9250) for 18 hrs, and then washed with PBS once, before adding fresh media containing 25 μM 2’-deoxycytidine (Sigma-Aldrich, D3897) for 9 hrs. After 9 hrs, 2 mM thymidine was added for 15 hrs. Cells were washed three times in PBS and released in fresh media containing 25 μM deoxycytidine for synchronous progression of the cell cycle from the G1/S transition.

### 2.3. siRNA transfection

HEK293ET or HeLa were seeded one day prior to transfection into DMEM + 10% (v/v) FBS, without antibiotics present. Twenty-four hours later, HEK293ET/HEK293T cells and HeLa were transfected using Lipofectamine^®^ 2000 or RNAiMAX, respectively (Thermo Fisher Scientific, Waltham, MA, USA). For one well from a 24-well plate 20 pmol of siRNA (HEK293ET) or 25 pmol (HeLa) were transfected in OptiMEM according to the manufacturer’s protocol. For U87 and U251 siRNA transfection, RNAiMAX was used according to the manufacturer’s protocol (Thermo Fisher Scientific).

All siRNAs were obtained from GE Dharmacon™ including On-TARGETplus Non-Targeting Pool siRNA (D-001810-10-05), On-TARGETplus HECTD1 Individual #06 siRNA (J-007188-06), #07 siRNA (J-007188-07), #08 siRNA (J-007188-08), #09 siRNA (J-007188-09) and On-TARGETplus HECTD1 SMARTpool siRNA (J-007188-00).

### 2.4. DNA transfection

HEK293ET and HEK293T cells were seeded one day prior to transfection into DMEM + 10% (v/v) FBS, without antibiotics present. Twenty-four hours later, the transfection mix was made by combining branched PEI (MW ~25,000) (Sigma-Aldrich, St Louis, MI, USA, 408727) and DNA at a ratio of 3:1, diluted into 150 mM NaCl. PEI and DNA (250 ng for one well of a 24-well) were individually diluted in 150 mM NaCl, left for 5 min before being combined and then incubated for 15 min at room temperature RT prior to addition to each well.

### 2.5. Propidium Iodide flow cytometry cell cycle analysis

HeLa cells were seeded at 300,000 cells per well and HEK293ET cells at 500,000 cells per well in a 6-well plate, prior to addition of the appropriate cell synchroniser. At each time point, samples were harvested in the following manner. The culture media was removed and saved in an Eppendorf, the cells were then rinsed with PBS, prior to addition of 300 μl of 0.05% Trypsin-EDTA (Thermo Fisher Scientific). Once the individual cells were fully detached, the media was added back to generate a single cell suspension. Cells were pelleted at 500 x g for 5 min, and the pellet was washed in PBS. Cells were fixed in 69% ethanol, 400 μl ice-cold PBS with 900 μl ice-cold 100% ethanol. Samples were then stored at 4°C for at least 2 hrs before staining. Samples were stained and data collected on the same day. Two μg/ml Propidium Iodide (PI) (Thermo Fisher Scientific, P3566) and 100 μg/ml RNase A (Thermo Fisher Scientific, EN0531) were diluted in 500 μl flow cytometry staining buffer (100 mM Tris pH 7.4, 150 mM NaCl, 1 mM CaCl2, 0.5 mM MgCl2, 0.1% Nonidet P-40). The samples were then incubated at 37°C for 30 min before being placed on ice in the dark, just before loading on the BD FACSCanto (BD Biosciences, San Jose, CA, USA). Data were analysed using BD FACSDIVA™ software V.8.0.1. For cell cycle analysis, histograms were gated at 50 PI-A to calculate the percentage of cells in G1 and 100 PI-A to calculate the % of cells in G2.

For flow cytometry analysis following SAC activation by Nocodazole, HEK293ET cells were first treated with either non-targeting siRNA (NT siRNA) or HECTD1 SMARTpool siRNA (HECTD1 SP) for at least 48hrs followed by Nocodazole treatment (18 hrs, 50 ng/μl). DMSO was used for the control experiment as indicated. Following treatment, cells were harvested and handled as described above.

### 2.6. Proliferation assay – trypan blue

HEK293ET and HEK293T cells were seeded at 60,000 cells per well in a Poly-L-Lysine coated 24-well plate (Corning Inc., Corning New York, USA, 3524). At each time point cells were trypsinised and resuspended in DMEM + 10% (v/v) FBS, before being mixed in a 1:1 ratio with Trypan Blue solution 0.4% (Thermo Fisher Scientific). Cells were then counted under a light microscope using a haemocytometer. Blue cells indicated dead cells. Counts were carried out in triplicate for each sample over three independent experiments.

### 2.7. Proliferation assay - CellTiter-Glo^®^ assay

HEK293T cells were seeded at 4,000 cells per well into a Poly-L-Lysine coated 96-well clear-bottomed white walled plate (Corning, 3903) in DMEM + 10% (v/v) without antibiotics. 48 hrs later cells were transfected with either empty vector (EV), full-length mouse HA-Hectd1^WT^, or HA-Hectd1^C2587G^ vector (Gift from Professor Irene Zohn, Children’s National Research Institute, USA), then left for 48 hrs before measuring the ATP content using CellTiter-Glo^®^ assay kit (Promega, Madison, WI, USA, G7570) [47]. The assay was carried out according to the manufacturer’s protocol for a 96-well format and measured using the GloMax Multi Plate Reader (Promega).

### 2.8. Immunofluorescence

For confocal imaging, cells were plated at an experiment specific density onto Poly-L-Lysine (mol wt 30,000 – 70,000) (Sigma-Aldrich, 9155) coated Nunc™ Thermanox™ 13 mm coverslips (Thermo Fisher Scientific, 1749500) in 12-well plates (Corning, CLS3513). Asynchronous or synchronous cells were then fixed with 4% (w/v) paraformaldehyde and stained. Cells were permeabilized in IF blocking buffer (PBS, 3% BSA, 0.1 Triton X100) for 5 min at RT. Primary and secondary antibodies were incubated in IF blocking buffer (PBS, 3% BSA) for 1 hr at RT each. Cells were washed twice with PBS and then counterstained with Hoechst 33342 (Thermo Fisher Scientific, H1399; 1 μg/ml in PBS), followed by two more washes in PBS. Coverslips were mounted with VectaShield Antifade Mounting Medium (Vector Laboratories, Burlingame, CA, USA; H-1000). Samples were then imaged on a LSM Meta 510 confocal microscope (Zeiss, Oberkochen, Germany). Primary antibodies included Anti-α-tubulin (Abcam, Ab7291, 1:1,000), Anti-HECTD1 (N-terminus) (Abcam, Ab101992, 1:200), Anti-HA (Roche, Switzerland; HA3F10, 1:1,000). Secondary included Alexa Fluor488 Goat anti-Mouse (Thermo Fisher Scientific’ A11029, 1:500), Alexa Fluor546 Donkey anti-Rabbit (Thermo Fisher Scientific; A10040, 1:500), Alexa Fluor456 Goat anti-Rat (Thermo Fisher Scientific; A11081, 1:500).

### 2.9. Live cell imaging

For live cell imaging, HEK293ET and HEK293T cells were seeded at 30,000 cells per ml onto Poly-L-Lysine (MW 30,000 - 70,000) (Sigma-Aldrich, 9155) coated plastic 8-well chamber slides (Ibidi IB-80826). Cells were left overnight in 300 μl/well DMEM + 10% (v/v) FBS, before exchanging the media the following day for Leibovitz’s L-15 media (no phenol red) (Thermo Fisher Scientific, 21083027), with 10% (v/v) FBS. Cells were then filmed over the course of hours at either 2, 3, or 5 min intervals using an Olympus IX81 microscope with a 40X oil immersion objective lens and Hammatsu ORCA-ET Camera for imaging at 37°C. Micro-Manager was used to acquire and manage the images [52]. Only the DIC channel was used for imaging. A binning of 2 was used for image acquisition, and images were acquired as a stacked TIFF format for downstream analysis. Images were then processed using ImageJ software [53]

### 2.10. Western blot analysis

Control asynchronous or synchronised cells were grown in an appropriately sized plate depending on the assay type, then lysed with either RIPA or Triton X-100 lysis buffer supplemented with Pierce™ Protease Inhibitor Mini Tablets (Thermo Fisher Scientific, 86665) and Pierce™ Phosphatase Mini Tablets (Thermo Fisher Scientific, 88667). Cell lysates were clarified by centrifugation and samples were denatured at 95°C for 5 min in NuPage 2X LDS/100 mM DTT sample buffer. Samples were run on NuPAGE^®^ Novex^®^ 4-12% Bis-Tris Protein Gels, 1.0 mm gels (Thermo Fisher Scientific, NP0322BOX) for 100 min at 140V in 1X NuPAGE^®^ MOPS SDS Running Buffer (Thermo Fisher Scientific, NP0001). Samples were run alongside the PageRuler™ Prestained Protein Ladder (Thermo Fisher Scientific, 26616). For immunoblotting, samples were transferred onto Whatman^®^ Westran^®^ PVDF membrane 0.45 μm using the Bio-Rad Mini Trans-Blot^®^ Wet Transfer System (Bio-Rad, Hercules, CA, USA), for 60 min at 100V in a Wet Transfer system. PVDF membranes were blocked in WB blocking buffer (3% BSA-PBS-Tween 0.1%) for 1 hr at RT. Primary antibodies were diluted in WB blocking buffer and incubated either over-night at 4°C or 1 hr at RT. Membranes were then washed in PBS-Tween (0.1%) and then incubated with species-specific horseradish-peroxidase (HRP)-conjugated secondary antibodies diluted in WB blocking buffer for 1hr at RT. Membranes were washed prior to detection with the Pierce™ ECL Western Blotting Substrate (Thermo Fisher Scientific, 32106). Chemiluminescence was detected using Fusion SL Chemiluminescence and Fluorescence Imager (Vilber Lourmat, France). Quantification of detected bands was carried out using ImageJ software [53].

Primary antibodies included: Anti-HECTD1 (N terminus) (Abcam, Ab101992, 1/2,500), Anti-Cyclin B1 (Santa Cruz Biotechnology, sc-245, 1/1,000), anti-phospho-Histone H3 (Ser28) (Abcam, Ab10543, 1/1,000), anti-β-actin (Sigma-Aldrich, A5441, 1/10,000), anti-CENPF (Abcam, Ab5, 1/1,000), anti-BUB3 (Abcam, Ab133699, 1/1,000), anti-BUBR1 (Abcam, Ab215351, 1/1,000), anti-MAD2 (Bethyl Laboratories, A300-310A, 1/1,000), anti-Plk (Santa Cruz Laboratory, F-8/sc-17783, 1/1,000), anti-GAPDH (Santa Cruz Laboratory, sc-25779, 1/2,000), phospho-Chk2 Thr68 (Cell Signaling Technology, #2661, 1/1,000), p21Waf1/Cip1 (Cell Signaling Technology, #2947) and PARP (Cell Signaling Technology, #9542, 1/1,000). Secondary antibodies included Goat anti-Mouse IgG HRP (Santa Cruz Biotechnology, sc-2054, 1/5,000), Goat anti-Rabbit IgG HRP (Santa Cruz Biotechnology, sc-2005, 1/5,000), Goat anti-Rat IgG HRP (Santa Cruz Biotechnology, sc-2006, 1/5,000), Rabbit anti-Goat IgG HRP (Thermo Fisher, 31402, 1/5,000).

### 2.11. GST pulldown

For the enrichment of ubiquitin chains, GST-TRABID NZF 1-3 was used while GST-TRABID NZF 1-3^TY>LV^ ubiquitin binding deficient mutant and/or GST were used as controls. All recombinant proteins were expressed in E. Coli BL21DE3(RIL) and purified a previously described [54]. For each 10 cm^3^ dish, 100 μg of trapping protein was used. 10 μg of GST tagged protein was conjugated to Pierce™ Glutathione Magnetic Agarose Beads (Thermo Fisher, 78602), for 1hr at RT. Cells in 10cm^3^ dishes were lysed in 500 μl of GST-TRABID NZF 1-3 IP Lysis Buffer (150 mM NaCl, 50 mm Tris pH 7.4, 5 mM DTT, 2 mM NEM, 10 mM iodoacetamide, 1% v/v Triton X-100)) for 20 min before being clarified by centrifugation (13,000 x g for 15 min at 4°C). 60 μl of lysate was collected for the input sample. 110 μl of lysate was added to 50μl of conjugated beads in 500 μl GST-TRABID 1-3 IP Pull-Down Buffer overnight at 4°C (150 mM NaCl, 50 mM Tris pH 7.4, 5 mM DTT, 2 mM NEM, 10 mM iodoacetamide, 100 μM ZnCl2, 0.1% (v/v) Nonidet P-40, 0.5 mg/ml BSA). Beads were then washed four times in GST-TRABID NZF 1-3 IP Wash Buffer (250 mM NaCl, 50 mM Tris pH 7.4, 5 mM DTT, 0.1% (v/v) Nonidet P-40) for 5 min, before the addition of 2X LDS sample buffer with 100 mM DTT.

### 2.12. Mass spectrometry analysis

HEK293T cells were synchronised in late G2 and mitosis as described above. Four μg of IgG or of a HECTD1 antibody (Ab101992) were coated onto magnetic Dynabeads^®^ magnetic beads and incubated with either lysates from G2 or M-phase synchronised cells for 1 hr at RT prior to washes, and denaturation in 2xLDS sample buffer/100 mM DTT at 95°C for 5 min. Samples were then loaded onto a 4-12% Bis-Tris SDS polyacrylamide gel which was then stained with Imperial Protein Stain (ThermoFisher Scientific). Gel lanes were sliced into of 1-2 mm^2^ pieces and following in-gel trypsin digestion followed by liquid chromatography tandem mass spectrometry (LC-MS/MS) using an Orbitrap Q Exactive (ThermoFisher Scientific) mass spectrometer coupled to a nano-ultra performance liquid chromatography (UPLC) system (Dionex, Sunnyvale, CA, USA) as described previously [36]. LC-MS/MS data were searched with the MASCOT program [55] against the UniPROT protein database. MS/MS data were validated using Scaffold (http://www.proteomesoftware.com).

### 2.13. Immunoprecipitation assays

For immunoprecipitation studies, endogenous HECTD1 was immunoprecipitated from either asynchronous HEK293ET cells, cells synchronised in G2 with RO3306, synchronised in G2 with RO3306 and released in mitosis (10 min and 30 min post-release from RO3306 block), or treated with Nocodazole (18 hrs, 50 ng/ml) to induce the SAC, as indicated. 4 μg of anti-HECTD1 antibody or normal Rabbit IgG was coupled to Dynabeads^®^ magnetic beads protein G (ThermoFisher Scientific, #10003D) for 1 hr at RT. Anti-HECTD1-coupled and IgG-coupled beads were washed 3x times in Triton Lysis Buffer prior to incubation with cell lysates. Cells were lysed on ice for 20 minutes in Triton Lysis Buffer supplemented with 1X Pierce™ Protease Inhibitor and 1X Pierce™ Phosphatase Inhibitor and the lysates was then cleared by centrifugation. HECTD1 or normal Rabbit IgG-coated beads were then incubated for 1 hr with cell lysates at RT on a rotating wheel. Magnetic beads from each condition were then washed 5x times in Triton Lysis Buffer and finally denatured at 95°C for 5 min in 2X LDS/100 mM DTT. Samples were then resolved on 4-12% SDS PAGE gels and analysed by immunoblotting. Typically, the input ran on a gel represented 1-5% of the total lysate [56].

### 2.14. Statistical analysis

All data were analysed using GraphPad software (GraphPad Prism, La Jolla, CA, USA), including mean, standard error values and statistical analysis. Standard error of the mean (S.E.M.) was calculated to quantify the precision of the mean and to compare differences in the means between conditions. S.E.M. was used to consider both the value of the standard deviation and the sample size. Statistical analysis was carried out either using a one-way ANOVA with a Dunnett’s post-test, or a paired student’s t-test as indicated. A one-way ANOVA with a Dunnett’s post-test was used when comparing a series of conditions to a single control condition. A paired student’s t-test was used when comparing two conditions. Relevant p-values are indicated where relevant by * p < 0.05, ** p < 0.01 and *** p < 0.001. A p-value of less than 0.05 was considered to be significant.

### 2.15. High-Content microscopy

10,000 cells were seeded into black-walled 96-well plates (Corning, 3340). For HEK293T cells, plates were also coated with 0.1 mg/mL poly-L-lysine (Sigma-Aldrich, P9155) for 30 minutes. After 24 hours, cells were incubated with 10 μM EdU (Invitrogen Click-iT™ EdU Alexa Fluor™ 488 HCS Assay, C10350) for 1 hour followed by PFA fixation and click-labelling according to manufacturer’s instructions. Following EdU labelling, cells were immunostained. Cells were blocked with 3% BSA in PBS for 1 hour at room temperature. Primary antibodies included phospho-Histone H3 (Ser28) (Abcam, Cambridge, UK; Ab10543, 1/200), phospho-γH2AX S139 (Cell Signaling Technology, Dancers, MA, USA; #9718; 1/200), phospho-Chk2 Thr68 (Cell Signaling Technology, #2661, 1/200). Secondary antibodies anti-Rabbit 555 (Ab150074, 1/500), anti-Rabbit 488 (Ab150073, 1/500) were from Abcam and anti-Rat 555 (A10522, 1/500) was from Invitrogen/Thermo Fisher Scientific. Antibodies were diluted in 3% BSA in PBS and incubated for 1 hour at room temperature. Nuclear counterstaining was done using 1X HCS NuclearMask™ for 30 minutes (Thermo Fisher Scientific). Plates were imaged using an IN Cell Analyzer 2000 (Cytiva, Marlborough, MA, USA), followed by image processing using CellProfiler v4.2.0 (https://cellprofiler.org) and data analysis using RStudio (R version 4.1.0) and GraphPad Prism v9.1.1. R Core Team (2021). R: A language and environment for statistical computing. R Foundation for Statistical Computing, Vienna, Austria. URL https://www.R-project.org/. GraphPad Prism version 9.1.1 for Windows, GraphPad Software, San Diego, California USA, www.graphpad.com.

## 3. Results

### 3.1. HECTD1 ubiquitin ligase activity contributes to cell proliferation

While previous studies have established roles for HECTD1 in Wnt signalling, transcription, cell migration and Base Excision Repair (BER), we made the serendipitous discovery that HECTD1-depleted cells grew more slowly than wild-type cells and followed up on this observation. We first used trypan blue dye exclusion to determine the effect of HECTD1 depletion on cell proliferation and found that transient HECTD1 siRNA KD in HEK293ET cells indeed decreased cell number, at day four post-transfection, compared to a non-targeting siRNA (Fig 1A & C). Similar data was obtained in HeLa cells and GBM cell lines U87 and U251, as well as in two independent HECTD1 KO HEK293T clonal cell lines (Fig 1D, E & Supp Fig 1A & B). This observed decrease in cell proliferation was not due to increased cell death as the percentage of viable cells remained similar for up to four days for the siRNA experiment, and 6 days for the KO cells (Supp Fig 1C-G). To validate this further, we used the CellEvent™ Caspase 3/7 Green detection reagent which indeed revealed that HECTD1 depletion does not increase caspase mediated cell death under basal conditions (Supp Fig 1H). We also used PARP cleavage as marker of caspase-mediated cell death but did no detect any PARP cleavage in wild-type nor in HECTD1-depleted HeLa or HEK293T cells (Fig 2H & data not shown). This suggests that the reduced cell proliferation observed upon HECTD1 depletion is not due to increase cell death.

**Figure 1:**
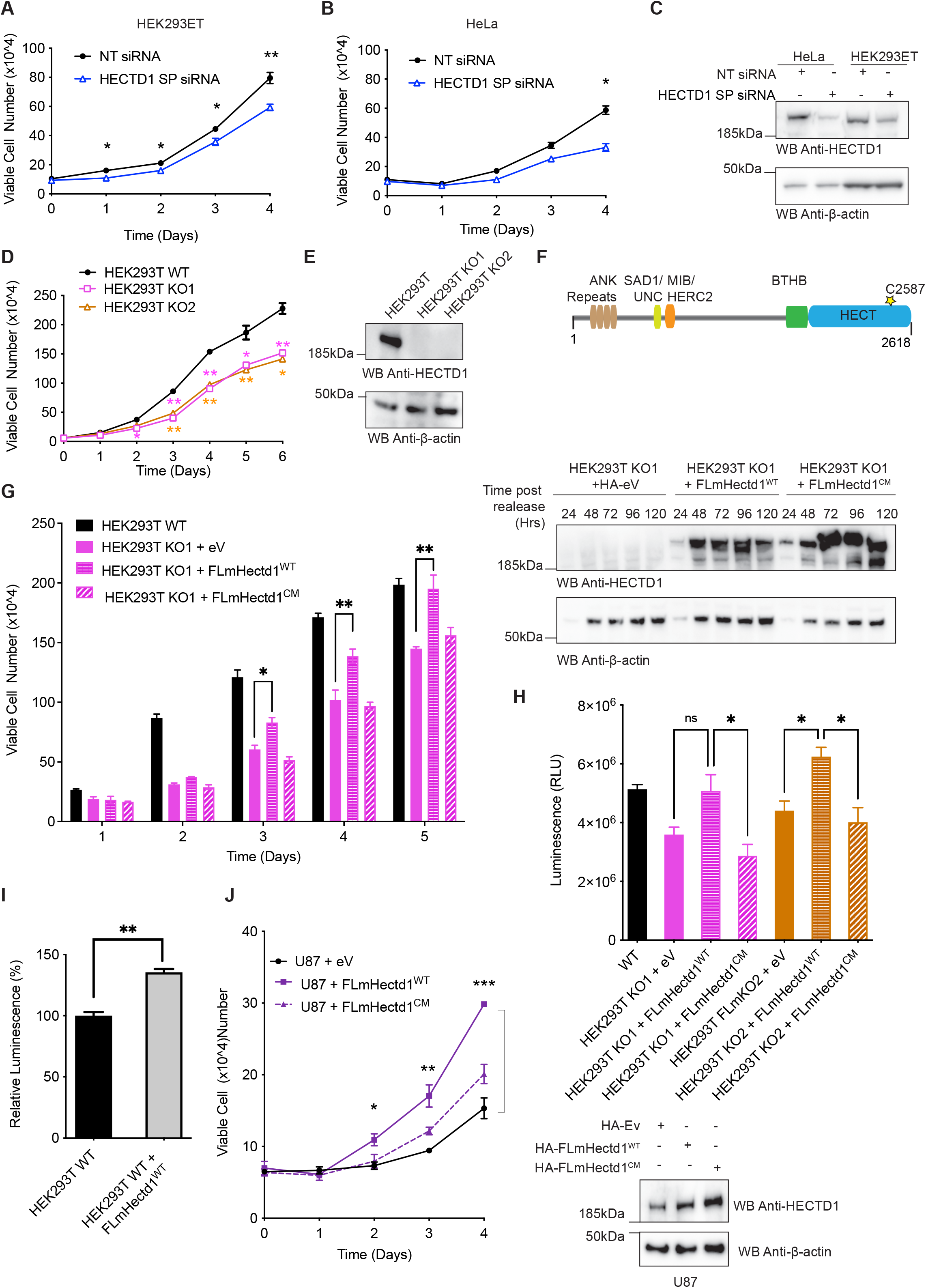
HECTD1 depletion reduces cell proliferation. **A-B)** The Effect of transient SIRNA knock down of HECTD1 on cell number was determined using trypan blue exclusion in HEK293ET (**A**) and HeLa (**B**). Data was plotted as mean with error bars that represent ±S.E.M., over three independent experiments (n=3) defined as three separate transfections, **p<0.01, *p<0.05 by paired student’s t-test. **C)** Immunoblot analysis showing HECTD1 transient knock down using 20 pmol SMARTpool (SP) S<RNA in HEK293ET and HeLa following 72 hrs incubation. HEK293ET and Hela cells were transfected with lipofectamine 2000 and RNAiMAX, respectively. Cells harvested at the indicated timepoint, lysed in RIPA buffer, and lysates were analysed on a 4-12% SDS PAGE followed by western blot analysis using an anti-HECTD1 antibody and an anti-β-actin antibody as loading control. **D)** Cell number was also quantified in HEK293T cells and two independent HECTD1 KO cell lines, KO1 and KO2. Data was plotted as mean with error bars that represent ±S.E.M., over three independent experiments (n=3) defined as three separate transfections, **p<0.01, *p<0.05 by paired student’s t-test. **E)** Immunoblot analysis showing the absence of HECTD1 in HECTD1 KO1 and KO2 cell lysates. **F)** Domain organisation of mouse Hectd1 highlighting HECTD1 catalytic cysteine (C2579 in human and C2587 in mouse). **G)** Rescue assay showing that re-expression in HEK293T KO1 of HA-FL-mHectd1 wild-type (WT), but not the catalytic mutant C2587G or an empty vector control, yields a similar number of cells to HEK293T WT cells. Data was plotted as mean with error bars that represent ±S.E.M., over three independent experiments (n=3) defined as three separate transfections, **p<0.01, *p<0.05 by a one-way ANOVA with Dunnett’s post-test. Right panel shows expression levels of HA-tagged constructs. **H)** A similar trend was also observed using the Cell-Titre-Glo^®^ Luminescence cell viability assay. The indicated samples were taken at 48 hrs post-transfection prior to analysis. Data plotted as mean with error bars that represent ±S.E.M., over three independent experiments (n=3) defined as three separate transfections, *p<0.05 by a one-way ANOVA with Dunnett’s post-test. **I)** HEK293T WT cells were transfected with HA-FL-mHectd1^WT^ or an HA-empty vector (ev) using PEI. 48 hours post-transfection cells were harvested, and cell proliferation measured (relative luminescence) with CellTiter-Glo^®^. Data is plotted as mean with error bars that represent ±S.E.M, over three independent experiments (n=3), **p<0.01 by a paired student’s t-test. **J)** Viable cell count (x10^4^) for U87 cells transfected with 250 ng either HA-empty vector (Ev), HA-FL-mHectd1^WT^, or HA-FL-mHectd1^C2587G (CM)^ using PEI. Samples were harvested at the indicated times post-transfection and counted using trypan blue to quantify the number of viable cells. Data plotted as mean with error bars that represent ±S.E.M., over three independent experiments (n=3). ***p<0.001, **p<0.01, *p<0.05 by one-way ANOVA with a Dunnett’s post-test. Right panel shows expression levels of HA-tagged constructs in U87.

**Figure 2.**
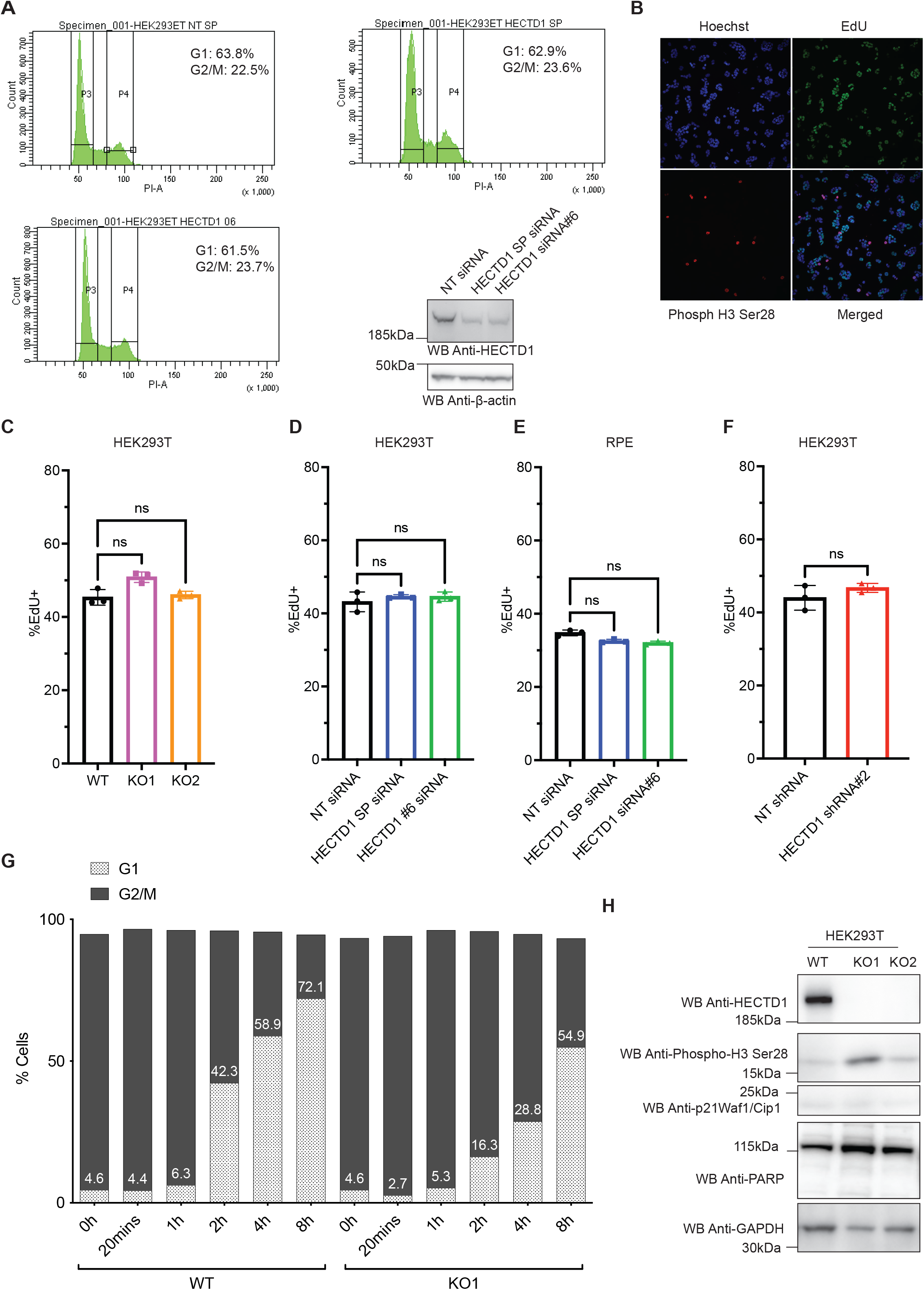
Cell cycle analysis of HECTD1-depleted cells. **A)** Flow cytometry analysis of the cell cycle in HEK293ET wild-type and HEK293ET cells treated for 48 hrs with either HECTD1 SMARTPpool (SP) siRNA or the individual SMARTpool HECTD1 siRNA #6. **B)** Representative images of HEK293T cells stained for EdU, phospho-Histone 3 (Ser28), Hoechst and imaged using an IN Cell Analyzer 2000 high-content microscope. Click-EdU^TM^ staining was used as a readout for cells in S-phase and quantified relative to the total number of cells in HEK293T WT, KO1 and KO2 **(C)**; in HEK293T **(D)** or hTERT-RPE cells **(E)** treated with HECTD1 SMARTpool siRNA or siRNA #6; or in individual NT-shRNA or HECTD1-shRNA clones **(F)**. Data plotted as mean with error bars that represent ±S.E.M., over three experiments (n=3 wells for each condition). Data analysed by unpaired t-test with Kruskal-Wallis. **G)** HEK293T WT or KO1 cells were synchronised in late G2 with 9 μM RO3306 for 20 hrs, and then released from block in full media. At each of the indicated timepoints, cells were fixed using 70% ethanol, and stained using 2 μg/ml PI, with 100 μg/ml RNase A, for 30 min at room temperature. Stained samples were then analysed immediately by flow cytometry. Gated population percentages are indicated on each graph. PI-A of 50 is equivalent to 2N (G1 population), and PI-A of 100 is equivalent to 4N (G2/M population). Graph showing the percentage of G1 and G2/M populations in HEK293T WT and KO1 cell lines at each time point post RO3306 release. **H)** Immunoblot analysis of RIPA lysates from HEK293T wild-type, HECTD1 KO1 and KO2 cell lysates. No change in PARP cleavage or p21Waf1/Cip1 levels was observed in HECTD1-depleted cells. In contrast, an increase in the levels of phospho-H3 (Ser28) was detected in both KO lines. GAPDH was used as loading control.

HECTD1 contains a conserved C-terminal HECT domain, composed of an N- and C-lobe, with the latter harbouring a conserved cysteine residue (C2579 in Human and C2587 in Mouse) which is essential for ubiquitin ligase activity (Fig 1F). To establish whether the observed decrease in cell proliferation was due to loss of HECTD1 ligase activity specifically, we carried out rescue assays using HA-tagged full-length mouse Hectd1 (FL-mHectd1^WT^) or an inactive version of this construct (FL-mHectd1^C2587G^) in HEK293T HECTD1 KO1 cells (Fig 1G). Re-expression of HA-FL-mHectd1^WT^ but not catalytic-dead HA-FL-mHectd1^C2587G^ recovered cell proliferation to a similar level to wild-type HEK293T cells. These observations were also validated using the CellTiter-Glo^®^ luciferase-based proliferation assay which showed a similar trend (Fig 1H). Furthermore, overexpression of HA-FL-mHectd1^WT^ but not the catalytic mutant resulted in a significant increase in cell proliferation as measured with the Cell-Titre-Glo^®^ assay in HEK293T or by trypan blue cell counting in U87 (Fig 1I & J, respectively). Together, these results indicate that the ubiquitin ligase activity of HECTD1 is important for cell proliferation.

### 3.2 Cell cycle analysis of HECTD1-depleted cells

To establish whether the observed decrease in cell proliferation was due to an effect on the cell cycle, we carried out flow cytometry analysis. Following siRNA-mediated knockdown of HECTD1 in HEK293ET, only a marginal decrease in the proportion of cells in G1 and a similar increase of cells in G2/M was observed (Fig 2A, Supp Fig 2A). Since S-phase is not reliably measured by flow cytometry, we used High-Content Microscopy (HCM) to quantify 5-ethynyl-2’-deoxyuridine (EdU) incorporation (Fig 2B). We observed no change in the proportion of cells in S-phase in HECTD1 KO cells, in HEK293T or hTERT-RPE cells treated with HECTD1 siRNA for 48 hrs, or in a HEK293T shRNA-HECTD1 clone (Fig 2 C-F). To expand on our observation that depletion of HECTD1 affects cell cycle progression, HEK293T WT and HECTD1 KO1 cells were synchronised in late G2 using 9 μM of RO3306 for 20 hrs (Supplementary Figure 2B-C) [57]. The progression of cells following release from G2 block was monitored by flow cytometry at the indicated time points (Fig 2G, Supplementary Fig 2D & E). As early as 2 hrs post R03306 release, when cells should have already exited M-phase, a higher proportion of cells was detected in G2/M in the HECTD1 KO1 cell line (79.5%), compared to control wild-type cells (53.7%). HECTD1 is required for Base Excision Repair (BER) and to account for the possibility that the increase in the G2/M population might reflect delay in another cell cycle stage due to DNA damage checkpoints and BER activation, we probed HECTD1 KO cell lysates for p21Waf1/Cip1 and phospho-Chk2 (Thr68) [38]. BER is mainly carried out during G1-phase, but it has been reported to take place throughout the cell cycle. Phospho-Chk2 (Thr68) and phospho-γH2AX are both upregulated by oxidizing agent H_2_O_2_ or the methylating agent Methyl MethaneSulfonate (MMS) (Supplementary Fig 3A & B) [58, 59]. However, we did not detect any change in p21Waf1/Cip1 or phospho-Chk2 (Thr68) levels by western blot analysis or in the number of cells positive for these markers by HCM in HECTD1-depleted cells (Fig 2H, Supplementary Figure 3C-D). However, we did detect increased phospho-H3 (Ser28) levels in HEK293T HECTD1 KO1 and KO2 (Fig 2H). This prompted us to use high-content microscopy to quantify phospho-H3 (Ser28)-positive cells but did not observe a statistically significant change between HECTD1 KO and wild-type cells (Supplementary Fig 4).

### 3.3 Phosho-H3 (Ser28) levels are increased in HECTD1-depleted cells

Phosphorylation of Histone H3 (Serine 28) has been used extensively as a reliable marker for early mitosis. Histone H3 is phosphorylated during prophase, and dephosphorylated upon anaphase exit [60]. Given our earlier finding that HECTD1 KO cells showed increased phospho-H3 (Ser28) levels, we quantified and further validated this data using transient siRNA knock down in HEK293T cells (Fig 3A & B). Phospho-H3 (Ser28) levels were also somewhat increased in hTERT-RPE cells treated with HECTD1 SMARTpool siRNA, in the absence of any detectable change in phospho-Chk2 (Thr68) (Figure 4B & Supplementary Fig 3E). Further analysis of HCM data confirmed this and revealed that at the population level, signal intensity of phospho-H3 (Ser28)-positive cells is also increased in HECTD1 siRNA-treated HEK293T cells compared to non-targeted siRNA treated cells (Fig 3C).

**Figure 3.**
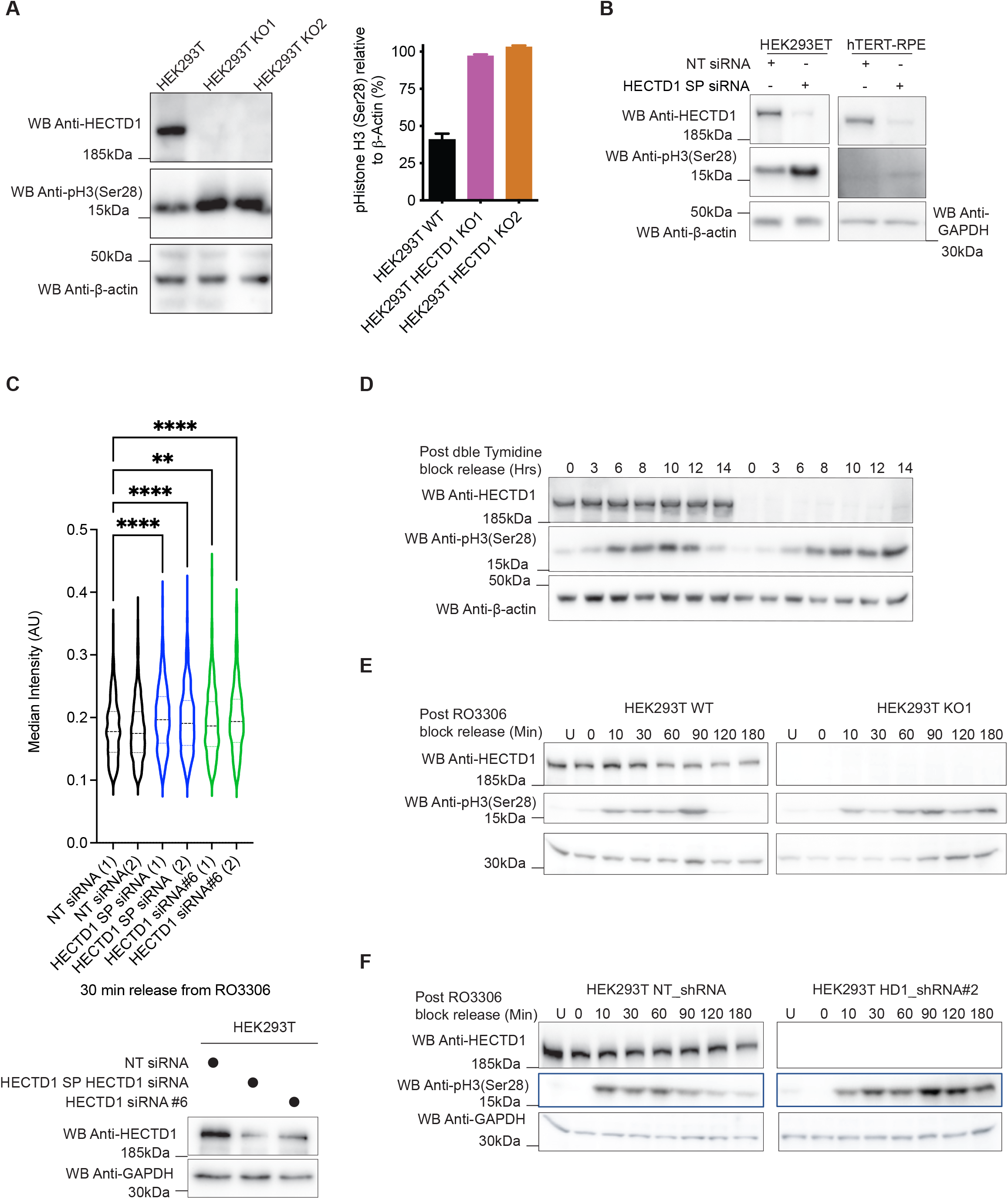
HECTD1 depletion increases phospho-H3 (Ser28) protein levels. **A)** Phospho-H3 (Ser28) levels were assessed in HEK293T wild-type and KO1 and 2 cell lysates, and changes were quantified and normalized to β-actin. Representative immunoblot analysis of HEK293T KO cells compared to WT control (n=2). **B)** A similar effect on Phospho-H3 (Ser28) levels was also observed following transient transfection with HECTD1 SMARTpool (SP) siRNA for 72 hrs in HEK293Tand 48 hrs in hTERT-RPE cells. **C)** Quantification of phospho-H3 (Ser28) median signal intensity obtained by high-content microscopy analysis of HEK293T cells treated with NT, HECTD1 SMARTpool or HECTD1 siRNA#2. Each condition was set up as two independent wells and data were analysed by One-way Anova. The lower panel shows HECTD1 levels in cells analysed by HCM for this experiment. β-actin or GAPDH was used as loading control. **D)** Western blot showing a phospho-H3 (Ser28) ‘tail’ in HECTD1-depleted cells compared to control (representative analysis of 2 independent experiments). Cells were treated with 2 mM Thymidine for 18 hrs, followed by a 9 hrs release. A second treatment with 2 mM Thymidine for 15 hrs was carried out, before releasing cells into full media. Samples were harvested at 0, 3, 6, 8, 10, 12, and 14 hrs post-release. Samples were collected at the indicated timepoints, lysed with RIPA and analysed by immunoblotting using anti-HECTD1, anti-phospho-H3 (Ser28) and anti-β-actin antibodies. **E-F)** Cell synchronisation assay using the CDK1 inhibitor RO3306. **E)** HEK293T WT or KO1 were synchronised in late G2 using 9 μM RO3306 for 20 hrs, and then released into full media to enter mitosis. Samples were harvested at the indicated timepoints post release, lysed in RIPA buffer, and probed for HECTD1, phospho-H3 (Ser28) and GAPDH. **F)** As in E) but using an individual NT-shRNA and a HECTD1-shRNA HEK293T stable cell line.

**Figure 4.**
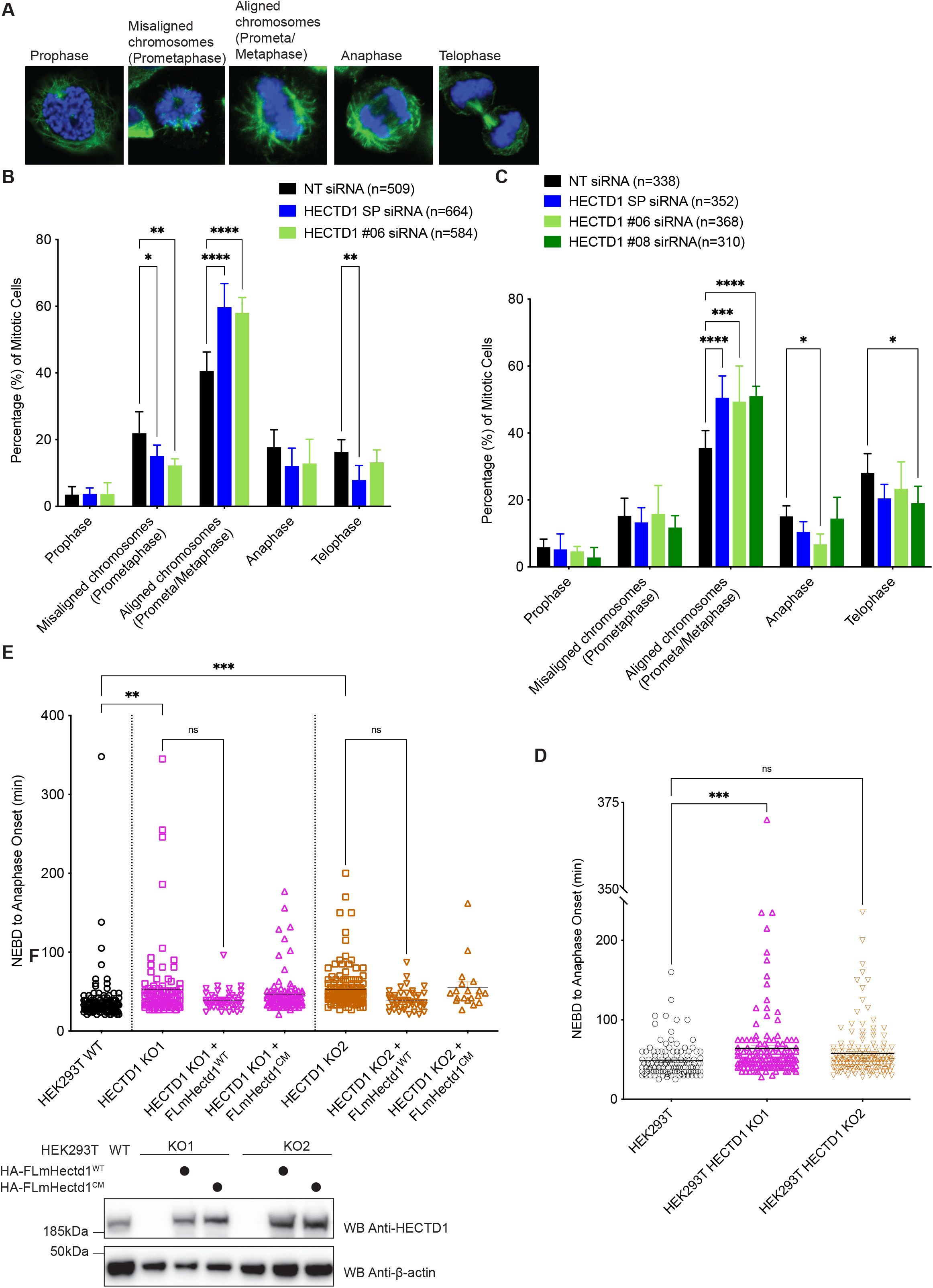
HECTD1 plays a role during mitotic progression. **A)** Confocal images of HEK293T cells at each mitotic stage which were used for B) and C). Cells were scored according to chromatin morphology based on Hoechst (blue) and α-tubulin (green) staining. Prometaphase refers to cells with misaligned chromosomes while Metaphase (i.e., Prometaphase/Metaphase) refers to cells with aligned chromosomes at the metaphase plate. **B)** HEK293ET cells and **C)** HeLa cells were scored according to chromatin morphology or spindle morphology, following 48 hrs transfection with Non-Targeting (NT) siRNA, HECTD1 SMARTpool (SP) siRNA, SMARTpool individual HECTD1 siRNA #06, or #08. Data plotted as mean with error bars that represent ±S.E.M., over 6 biological repeats (individual transfections). **p<0.01, ***p<0.001, and ****p<0.0001 using a one-way ANOVA with a Dunnett’s post-test. Each HECTD1 siRNA condition was significant when compared to the corresponding NT siRNA control. **D)** The duration of NEBD to anaphase onset was timed in asynchronous HEK293T WT and KO1 and KO2 cell lines. Vertical scatter plot showing the time taken for individual cells to progress from NEBD to anaphase onset. Error bars represent ±S.E.M., ***p<0.001, using a one-way ANOVA with a Dunnett’s post-test. Number of cells filmed are as follows, WT = 116, KO1 = 136, and KO2 = 161, filmed over 4 independent experiments. **E)** Vertical scatter plot showing the time taken (min) for each cell to progress from NEBD to anaphase onset in HEK293T KO1 cells transfected for 48 hrs with either HA-FL-mHectd1^WT^, or HA-FL-mHectd1^C2587G (CM)^. Lower panel shows expression levels of HA-tagged constructs.

To further explore the effect of HECTD1 depletion on mitotic entry and exit, we synchronised HEK293T WT and KO cells at the G1/S checkpoint using double thymidine block and released cells from this block for the indicated time. In this synchronisation experiment a signal for phospho-H3 (Ser28) was retained for longer in HECTD1 KO1 compared to wild-type HEK293T cells (Fig 3D). Although double thymidine block is one of the most used synchronisers for cell cycle studies, this treatment can also trigger DNA damage which may impact on cell cycle progression. Alternative compounds, including the Cdk4/6 inhibitor Palbociclib, have shown various efficacy at synchronising different cell lines [61]. Since our data argues that HECTD1 depletion increases phospho-H3 (Ser28) levels, we asked whether progression through M-phase, following release from RO3306-mediated late G2 block, might be impacted [57, 62]. Again, we found that phospho-H3 (Ser28) was retained for longer following release from RO3306-induced G2 block in the HECTD1 KO1 cell line as well as the HEK293T HECTD1 shRNA#2 stable cell line compared to controls (Fig 3E & F, respectively). This data suggests that HECTD1 contributes to cell cycle progression through a yet-to-be identified mechanism in mitosis.

### 3.4 HECTD1-depleted cells accumulate during mitosis

In order to further explore the impact of HECTD1 depletion on mitosis, we quantified mitotic phenotypes from confocal microscopy images (Fig 4). Mitotic cells were scored as prophase, misaligned chromosomes (prometaphase), aligned chromosomes (prometaphase/metaphase), anaphase and telophase, based on chromatin and microtubule arrangement (Fig 4A) [63]. HECTD1 was transiently depleted in HEK293ET and HeLa cells and stained with anti-α-tubulin antibody and Hoechst. In addition to using the HECTD1 SMARTpool we also tested the individual siRNAs #06, #07, #08 and #09 in HEK293ET and HeLa cells (Supplementary Fig 2A & F, respectively). In HEK293ET, 40% of mitotic cells treated with a non-targeting (NT) siRNA showed aligned chromosomes, compared to 57.7% for HECTD1-depleted cells using siRNA #06, and 58.1% for the HECTD1 SMARTpool (SP)-treated cells (Fig 4B). We observed a similar effect in HeLa cells where 35.7% of cells in the NT siRNA condition were scored as aligned chromosomes, compared to 50.4% for SMARTpool-siRNA, 48.8% for siRNA #06, and 51.7% for siRNA #08 (Fig 4C). HECTD1-depleted HEK293ET and HeLa cells showed an 18% and 15% increase in the number of cells with aligned chromosomes (i.e., apparent metaphase plate), respectively. Despite the accumulation of mitotic cells following HECTD1 knockdown, we did not observe any obvious or significant effect on mitotic spindles, suggesting that the observed enrichment of HECTD1 depleted cells in metaphase does not translate into mitotic defects in most cells (Supplementary Figure 5A-C). Although we did detect a slight increase in multinucleated cells and multilobed nuclei in HECTD1-depleted cells, this did not reach statistical significance (Supplementary Figure 5D-F).

### 3.5 Loss of HECTD1 ligase activity slows down mitosis

To further establish a novel role for HECTD1 in mitotic progression, we next used time-lapse microscopy to measure the time taken to progress from Nuclear Envelope Breakdown (NEBD) to anaphase onset as a readout for mitotic progression. Examination of the time-lapse image series of mitotic cells suggested that some cells depleted of HECTD1 showed chromosomes aligned along the equatorial plane for longer than wild-type cells before anaphase onset, and we therefore set out to quantify this change (Supplementary Fig 6A). Time measurement of NEBD to anaphase onset revealed that HEK293T_HECTD1_KO1 cells showed a significant mean delay of around 15.5 min and KO2 cells a mean delay of around 9 min compared to the HEK293T control cells (Fig 4D & Supplementary Fig 6B). Time-lapse microscopy of HEK293ET cells transiently transfected with HECTD1 SMARTpool or the indicated individual siRNAs also gave a similar trend (Supplementary Fig 6C & D). At 72 hrs post-siRNA transfection, HECTD1 SMARTpool showed a mean delay of 21.6 min, and HECTD1 #06 siRNA a significant mean delay of 39.3 min compared to the NT siRNA control. Although siRNA #8 showed no mean delay, individual cells still showed a trend towards delayed anaphase onset. Overall, these results suggest that the transient knockdown or genetic knockout of HECTD1 results in a mitotic delay of around 15-30 min on average, representing a 30-60 % increase in the duration of mitosis in HEK293T and HEK293ET cells, respectively, which was around 50 min for these cells. Together, these data suggest that some HECTD1-depleted cells progress more slowly from NEBD to anaphase onset and is in line with the observed increase phospho-H3 (Ser28) levels, and the increase in the proportion cells with aligned chromosomes.

To ascertain the role of HECTD1 ligase activity in this phenotype, we carried out rescue assays. NEBD to anaphase onset was measured in asynchronous HEK293T KO1 cells transfected with either HA-FL-mouse Hectd1^WT^ or HA-FL-mouse Hectd1^C2587G^ (Fig 4E & Supplementary Fig 6E). Re-expression of HA-FL-mHectd1^WT^ but not HA-FL-mHectd1^C2587G^ rescued the mitotic delay phenotype in both HECTD1 KO cell lines (Fig 4E). HEK293T wild-type cells took on average 37 min to progress from NEBD to anaphase onset. In comparison, both HECTD1 KO1 and KO2 cells showed an increase to 53 min on average. Importantly, HA-FL-mHectd1^WT^ rescued this delay by around 14 min on average. Taken together, this suggests that HECTD1 ubiquitin ligase activity can rescue the reduced cell proliferation phenotype and the NEBD to anaphase onset delay phenotype observed in HECTD1-depleted cells.

### 3.6 HECTD1 mitotic interactors

The Spindle Assembly Checkpoint (SAC) occurs during mitosis and prevents anaphase onset until each kinetochore is attached to the mitotic spindle, ensuring proper chromosome segregation [64]. This mechanism represents a possible explanation for the observed delay in HECTD1-depleted cells. To test this hypothesis, we explored whether HECTD1 depletion might have a stronger phenotype upon SAC activation. HEK293ET cells treated with either a non-targeting (NT) or with HECTD1 SMARTpool siRNA for 48hrs, followed by addition of DMSO or the potent activator of the SAC, Nocodazole (Fig 5A & B). Following this treatment, flow cytometry was carried out and cell cycle profiles analysed (Fig 5C & D). HECTD1 depletion reduced the proportion of cells in G2/M by over 8% upon SAC activation. This data suggests that HECTD1-depleted cells, under conditions with activate the SAC, are less efficient at arresting in mitosis.

**Figure 5.**
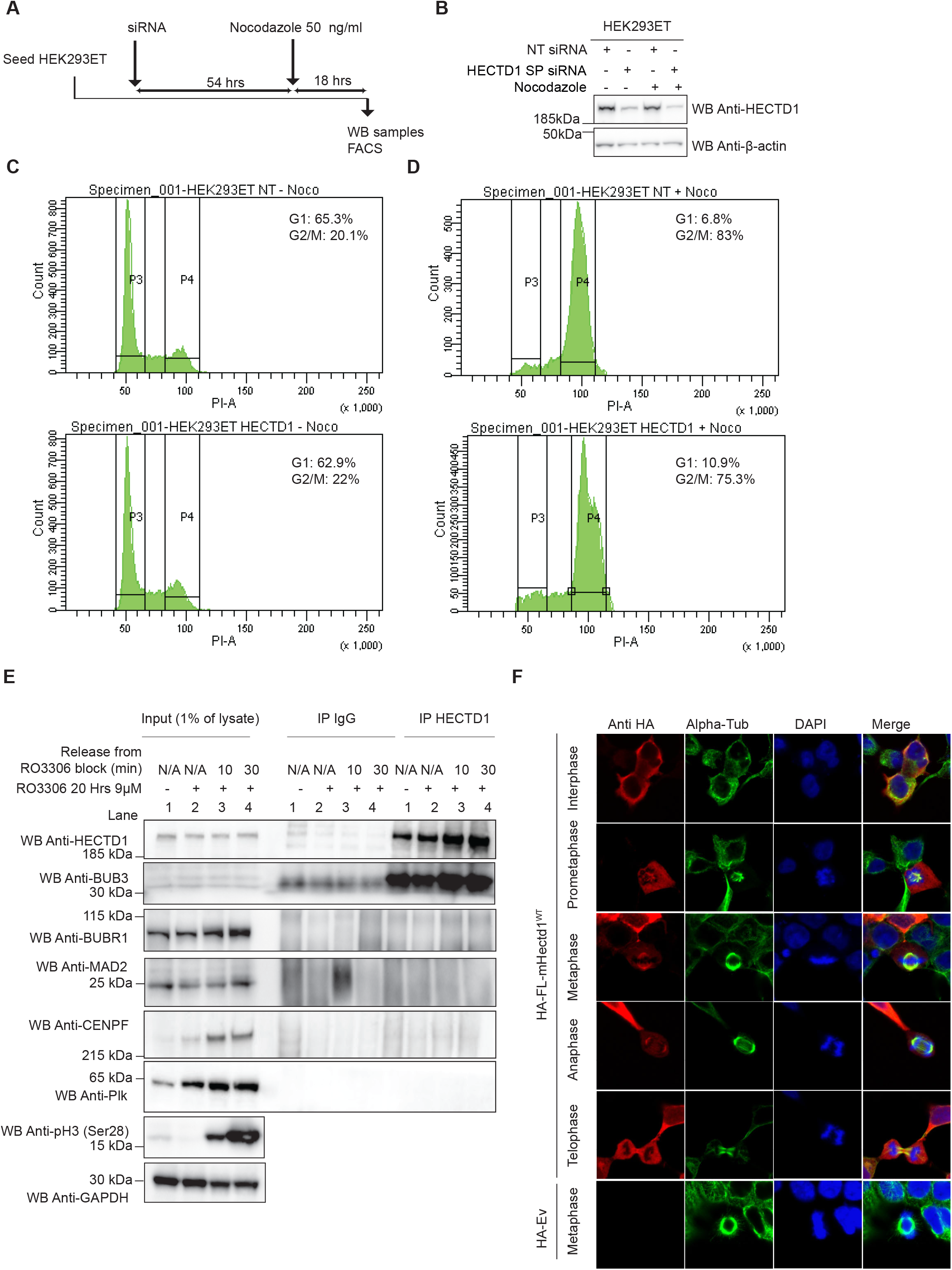
HECTD1 contributes to SAC activation. **A)** Experimental setup to test the effect of HECTD1 depletion on SAC activity. **B)** Immunoblot analysis showing HECTD1 levels following 48 hrs treatment with HECTD1 SMARTpool (SP) siRNA in HEK293ET cells. β-actin was used as loading control. **C-D)** Flow cytometry analysis of the cell cycle in HEK293ET treated for 48 hrs with NT or HECTD1 SP siRNA prior to addition of DMSO (**C**) or Nocodazole (50 ng/ml for 18 hrs) (**D**), as shown in A). **E**) Immunoprecipitation assay of endogenous HECTD1 in HEK293T cells showing interaction with endogenous BUB3, but not MAD2, BUBR1, CENPF or Plk. Normal Rabbit IgG was used for control IP. Note that the same results were obtained whether cells were asynchronous (Lane 1), synchronised in late G2 with RO3306 (Lane 2), or released from RO3306 block into mitosis for 10 min (Lane 3) or 30 min (Lane 4). Phospho-H3 (Ser28) was used to show the effective synchronisation using R03306, while β-actin was used as loading control. Input (1% of lysate) and IP materials were run onto the same gel. **F)** Representative confocal images of HA-FL-mHectd1^WT^ transiently expressed in HEK293ET cells. Cells were transfected with 500 ng of HA-mHectd1^WT^ or HA-tagged empty vector (EV) using PEI, in a 12-well format. 24 hrs post-transfection cells were fixed using 4% PFA, then stained with anti-HA (red), anti-α-tubulin (green), counter stained with Hoechst (blue) and mounted with VectaShield mounting media. Images were taken using an LSM Meta 510 Confocal Microscope. Scale bar represents 10 μm. Images were taken of cells in prometaphase, metaphase, anaphase, and telophase.

The time-resolved interactome of cyclins has revealed important insights with regards to the plasticity of cyclins complexes during cell cycle progression [65]. To try and further explore the mechanisms underlying HECTD1 putative function during mitosis, we carried out a proof-of-principle proteomics experiments to identify HECTD1 candidate interactors (Supplementary Fig 7B-D). HEK293T cells were synchronised in late G2 using RO3306 and IgG-coated or HECTD1 antibody-coated Dynabeads^®^ magnetic beads were used in immunoprecipitation (IP) experiments (Supplementary Fig 7B). We also determined the HECTD1 interactome 20 minutes post-RO3306 release to get insights on HECTD1 mitotic interactome. Pull-down materials from either the IgG control IP or the endogenous HECTD1 IP were then analysed by LC-MS/MS on an Orbitrap velos (Supplementary Fig 7C). From a list of 849 proteins that were identified (at least 2 unique peptides but 0 peptide in the IgG IP), we specifically searched for cell cycle-related proteins and found several candidate interactors of endogenous HECTD1 (Supplementary Fig 7D). We also identified distinct subsets of proteins enriched in the HECTD1 interactome in late G2 vs M-phase indicating that the HECTD1 interactome is dynamic during the cell cycle. Interestingly, we identified the Mitotic Checkpoint Complex protein BUB3 as a novel candidate HECTD1 interactor. BUB3 works as part of a multiprotein complex together with BUBR1, CDC20 and MAD2 to regulate the SAC [66–69]. The Mitotic Checkpoint Complex forms at unattached kinetochores, sequestering and inhibiting the APC/c ubiquitin ligase complex and preventing anaphase onset. Since flow cytometry experiments indicated a putative novel role for HECTD1 in full activation of the SAC, we carried out IP experiments which confirmed that endogenous HECTD1 indeed interacts with BUB3, but not with MAD2 or BUBR1 (Fig 5E). The endogenous HECTD1-BUB3 interaction could be detected in cells irrespective of whether cells were synchronised in late G2, M-phase or asynchronous. This suggests that HECTD1 interaction with BUB3 is constitutive and not induced by SAC activation. Although CENPF was also found in our proteomics list, we could not validate this interaction using our IP conditions. Polo-like kinase (Plk) served as negative control since we did not find it in our list of candidate interactors. To further explore the interaction between HECTD1 and BUB3, we next explored whether this interaction might be enriched in HEK293T cells under conditions where SAC was activated (i.e., Nocodazole treatment 50 ng/ml, 18 hrs) (Supplementary Figure 7E). However, this was not the case, suggesting that a pool of HECTD1 might be constitutively associated with BUB3 independently of the status of the SAC.

Since the commercially available HECTD1 antibodies which we tested produced either weak or non-specific signals by IF (not shown), we investigated the localization of HA-FL-mHectd1^WT^ or HA-FL-mHectd1^C2587G^ in transiently transfected cells (Fig 5F). Confocal microscopy revealed that in interphase cells, including in HEK293T and U87 cells, HA-FL-mHectd1 is primarily cytosolic (not shown), while it is found at mitotic spindles in cells undergoing mitosis, and this is independent of the ligase activity (Not shown).

### 3.7 TRABID NZF traps polyubiquitin in each phase of the cell cycle

The deubiquitylase TRABID/ZRANB1 preferentially recognizes and processes K29 and K33 linkages over any other linkage types including K63-linked chains [54, 70, 71]. TRABIDs’ three Npl14 zinc finger (NZF) domains act as ubiquitin binding domain, with NZF1 largely responsible for this linkage-specific binding [72–74]. TRABID has emerged as a unique reagent to capture and study atypical ubiquitin chains [54, 70–73]. Our recent work provided compelling evidence that HECTD1 catalytic HECT domain, preferentially assembles branched K29/K48-linked chains [36]. We next used TRABID NZF domain as the best available reagent to try and identify K29-linked chains during the cell cycle (Fig 6). First, we validated the interaction between TRABID and HECTD1 using GST-tagged TRABID NZF 1-3, in line with our previous work (Fig 6A & B). Treatment with the proteasomal inhibitor MG132 yielded more ubiquitin species as suggested by the increase in ubiquitin signal, further indicating that some of the ubiquitin chains captured by TRABID NZFs are associated with proteasomal degradation (Fig 6B) [75]. We confirmed that the signal indicative of interaction between the GST-TRABID NZF 1-3 bait and endogenous HECTD1 was lost in lysates from CRISPR/Cas9 HECTD1 KO cells (Fig 6C, lane 5 vs. 2). In line with our previous data, the TRABID-HECTD1 interaction was dependent on the ubiquitin binding property of TRABID NZF 1-3, as the TY>LV mutation in each of these UBDs abrogated binding to endogenous HECTD1 in our pull-down assay (Fig 6C, lane 3 vs 2) [36, 71]. Some polyubiquitinated species could still be detected in the GST-TRABID NZF 1-3 pull down with lysates from HECTD1 KO1 HEK293T cells (Fig 6C, lane 5 vs. 2). This indicates that although HECTD1 assembles a large proportion of the polyubiquitinated species trapped by TRABID NZF 1-3, some polyubiquitinated species are likely contributed by other E3s, most likely UBE3C and TRIP12 given these E3s also assemble K29-linked polyubiquitin or perhaps E3s assembling K33-linked chains [30, 34, 72, 73, 75].

**Figure 6.**
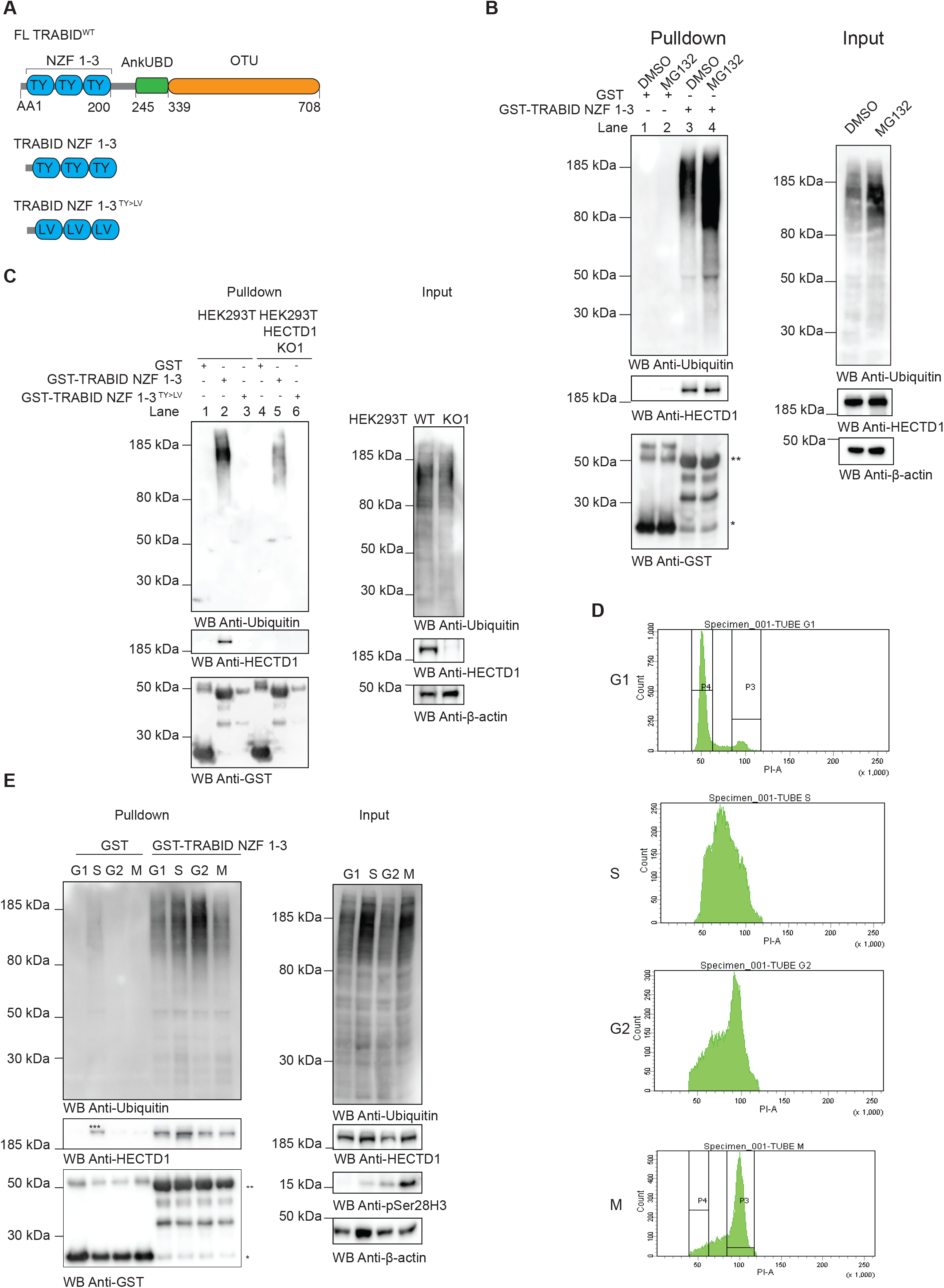
TRABID NZF 1-3 traps ubiquitin chains during cell cycle progression. **A)** Domain organisation of TRABID showing the AA1-200 region which contains three Npl14 UBDs. GST alone, GST-tagged TRABID NZF 1-3 and the ubiquitin binding deficient TRABID NZF 1-3 TY-LV were produced in E. coli used as baits in pull down experiments. **B)** GST alone or GST-TRABID NZF 1-3 were used as bait in pulldown experiments with cell lysates from asynchronous HEK293T WT cells treated with DMSO or 10 μM MG132 for 6 hrs prior to cell lysis. Following pull down with Pierce™ Glutathione magnetics agarose beads, samples were resolved on 4-12% SDS PAGE, transferred onto PVDF, and probed with anti-Ubiquitin or anti-HECTD1 antibodies. Anti-GST was used as loading control for the baits and β-actin was used as a loading control for lysates. Star (*) indicates GST and (**) indicates GST-TRABID NZF 1-3. **C)** GST-TRABID NZF 1-3 traps ubiquitin and its interaction with HECTD1 requires functional ubiquitin binding domains. Pulldowns and Input were carried out as in B) using lysates from HEK293T WT and HEK293T KO1 mutant cells. **D-E)** Pull down assay to determine the ability of GST-TRABID NZF 1-3 to pull down endogenous ubiquitin from synchronised HEK293T WT. HEK293T WT cells were synchronised using 4 μg/ml Aphidicolin to enrich for G1 phase cells, 2 mM Thymidine with a 2 hrs release (double thymidine block) to synchronise cells in S phase, 9 μM RO3306 for 20 hrs to obtain G2 and following 20 minutes release from RO3306 to obtain M-phase cells. **D)** Cell cycle analysis by flow cytometry showing. Histograms for each cell cycle phase is shown. PI-A of 50 is equivalent to 2N (G1 population), and PI-A of 100 is equivalent to 4N (G2/M population). **E)** Immunoblot analysis of pulldowns carried out using lysates from synchronised cell populations, using GST or GST-TRABID NZF 1-3 as baits. Anti-phospho Histone H3 (Ser28) and anti-β-Actin were used as control for M-phase and loading control, respectively. Note that the S-phase samples was overloaded compared to the other samples. * Represents GST, ** GST-TRABID NZF 1-3, and *** reflects uneven loading of the lysate obtained from the S-phase synchronisation.

Ubiquitin chains have shown some degree of specificity during cell cycle progression, with K48-linked ubiquitin chains being predominant in S-phase while K11-linked ubiquitin chains so far appear the main signal for UPS-mediated degradation during mitosis [15]. To further explore whether other atypical ubiquitin chains might be found in specific stages of the cell cycle, we used GST-TRABID NZF 1-3 to enrich for ubiquitin chains in lysates obtained from HEK293T cells synchronised in G1, S, G2 and M phases (Fig 6D & E). This revealed that the TRABID-HECTD1 interaction, as well as the polyubiquitinated species recognized by TRABID NZF 1-3, can be captured in all stages of the cell cycle, including mitosis. Taken together, our data raise questions with regards to the putative function of the chains assembled by HECTD1 in mitosis but also in other cell cycle stages.

## 4. Discussion

This study was initiated through the fortuitus observation that HECTD1 depletion led to a reduction in cell number, and we determined this was dependent upon the ubiquitin ligase activity of HECTD1. To further elucidate the cellular mechanisms involved, we investigated the impact of loss of HECTD1 on cell cycle progression. Interestingly, a recent study reported that HECTD1 depletion reduces protein synthesis and cell cycle kinetics during Haematopoietic Stem Cells (HSCs) regeneration [76]. Similarly, to this study, we found that HECTD1 depletion did not affect cell viability. This study also showed that HECTD1 depletion reduced HSCs in S-phase under stress conditions. However, under basal (i.e., non-stress) conditions, we observe no change in the number of cells in S-phase, which indicates that HECTD1 has different effects depending on whether cells are grown under basal or under stress conditions. Flow cytometry revealed only a small increase in the G2/M population upon HECTD1 transient siRNA depletion but we did observe an increase in the levels of the mitotic marker phospho-H3 (Ser28) in HECTD1-depleted cells.

Our data suggests a new and so far, unreported direct effect of HECTD1 depletion on the cell cycle, specifically during mitosis. In support of this, immunoblotting of synchronised cells, revealed that lysates from HECTD1-depleted cells retained the mitotic marker phospho-H3 (Ser28) for longer compared to wild-type cells. HECTD1 contributes to base excision repair (BER) and HECTD1 depletion can lead to deficiencies in DNA repair and decreased cell survival in clonogenicity assays [38]. The proposed mechanism appears to be mediated through the ubiquitylation of histones although the type of ubiquitin chains involved remain to be determined. In our assays however, carried out under basal/unchallenged conditions, we did not observe any change in the levels of DNA damage markers p21Waf1/Cip1, phospho-γH2AX or the BER marker phospho-Chk2 (Thr68) between HECTD1-depleted and control cells. This suggested that the effects we observed on mitosis is unlikely to be due to knock-on effects or repair mechanisms taking place at other stages of the cell cycle. To confirm this, we used time-lapse imaging of NEBD to anaphase onset in asynchronous cells as a readout for progression through mitosis. This revealed that HECTD1-depleted cells were indeed slower to progress through mitosis. Importantly, our rescue experiments also indicated that this phenotype could be rescued upon re-expression of full-length wild-type but not the catalytic inactive version of Hectd1. Delay in mitosis, measured by timing NEBD to anaphase onset, can be indicative of the SAC remaining active for longer. This leads to APC/c inhibition and delayed anaphase onset. In line with this hypothesis, single cell microscopy analysis revealed an enrichment of cells with aligned chromosomes upon transient siRNA knockdown of HECTD1. We also found that Nocodazole-mediated SAC activation was less efficient in HECTD1-depleted cells compared to control cells, further hinting at a possible role for HECTD1 in the SAC-associated regulation of mitosis. HECTD1 is not the first HECT E3 shown to participate in SAC-activation, with SMURF2 and EDD both previously implicated [27, 77]. This raises important questions including how the activity of different E3 ubiquitin ligases is coordinated during SAC activation.

Our pilot HECTD1 interactome study identified several candidate proteins relevant to the cell cycle. Aurora kinase B and other cell cycle proteins have been identified as candidate HECTD1 interactors in human Embryonic Stem Cells [78]. However, whether and how HECTD1 regulates the Chromosomal Passenger Complex remains to be determined. Using proteomics analysis, we identified BUB3 as a novel candidate HECTD1 interactor and validated that endogenous HECTD1 does indeed interact with BUB3 but not with BUBR1 or MAD2, two other components of the Mitotic Checkpoint Complex [79]. The HECTD1-BUB3 interaction is likely constitutive since it takes place in asynchronous cells, cells synchronised in late G2 or in M-phase and it is independent of SAC activation. Although HECTD1 depletion appears to reduce the activity of the SAC, protein levels of BUB3, BUBR1 or MAD2 do not appear to be affected. The RepID-CRL4 ubiquitin ligase complex regulates BUB3 levels during mitosis and it will be interesting to further evaluate the functional consequences of HECTD1 interaction with BUB3 [81]. Furthermore, Tandem Mass Tag (TMT)-Mass Spectrometry combined with proximity labelling would also provide novel insights on the role of HECTD1 in mitosis [65, 80].

Using TRABID NZF ubiquitin binding domain as bait, we provide proof-of-principle that ubiquitin chains likely containing K29-linkages can be trapped in different cell cycle stages. Although affimers specific for K33 and K6 ubiquitin linkages have been available for a few years, it is only during the preparation of this manuscript that a K29-specific affimer was reported [50]. Excitingly this study revealed the enrichment of K29-linked ubiquitin in stress response and cell cycle regulation. Reducing K29-linked ubiquitin through overexpression of TRABID, which cleaves these linkages, led to cell cycle arrest in G1/S phase. This data which implicates K29-linked ubiquitylation during cell cycle progression, also revealed the presence of K29 linkages in mitotic cells, specifically at the midbody during telophase. In line with our data, HECTD1 and BUB3 were identified from pulldown and proteomics analysis using the K29-specific affimer [50]. This exciting new reagent now enables further studies to explore and the function of ubiquitin chains containing K29 linkages in various cellular context including during cell cycle progression.

Interestingly, we also found that HECTD1 ubiquitin ligase activity enhances cell proliferation in glioblastoma cells and according to two Oncomine datasets, *HECTD1* mRNA is found overexpressed in some GBM samples (Supplementary Figure 8A). Although we are yet to establish the underlying mechanisms in these cancer cells, recent mechanistic studies have proposed that USP15 stabilizes HECTD1 levels, which may inhibit Wnt pathway activity and reduce GBM growth [82]. However, in the absence of rescue experiments with HECTD1, it remains unclear whether the HECTD1 ligase activity is in fact implicated. Furthermore, GBM exhibit high levels of heterogeneity and it will be important to dissect HECTD1 function in GBM subpopulations to provide a deeper mechanistic understanding. In TCGA datasets, high HECTD1 expression is found associated with decreased survival in the mesenchymal GBM subtype and in pancreatic adenocarcinoma (Supplementary Figure 8B and C) [83]. It will be important to further explore the role of HECTD1 in these cancers and evaluate its potential as therapeutic target [84].

## 5. Conclusions

Protein ubiquitylation is a major regulator of cell cycle progression including cell division. The APC/c ubiquitin ligase complex drives the degradation of mitotic cyclins through branched K11/K48-linked ubiquitin chains and this is key to mediate anaphase onset and trigger the end of mitosis. Yet, whether and how other E3 ubiquitin ligases and ubiquitin signals participate during the cell cycle, including in mitosis, remains less well understood. Our findings show that HECTD1 regulates cell proliferation through an effect on mitosis. Although the exact mechanisms taking place will need to be further explored, we identified BUB3 as novel HECTD1 interactor. Together with functional assays, our data suggest that HECTD1 activity is required for full SAC activation. Our previous work established that the catalytic HECT domain of HECTD1 preferentially assembles branched K29/K48 ubiquitin chains. Since homotypic K63 and K48-linked chains have also been proposed as possible signals assembled by HECTD1 ligase activity, it will be important to establish which is relevant during mitosis. Tandem Ubiquitin Binding Entities and the recently developed ubiquitin K29-specific affimer offer new and exciting opportunities to address these important questions. HECT E3 ligases have been linked with various aspects of tumorigenesis, and future work should also aim to evaluate HECTD1 in cancer, including its potential as a therapeutic target.

## 6. Author contribution

JDFL formulated the hypothesis, conceived, and managed the project, supervised, and trained students. CL trained and supervised NV for the live cell imaging experiments which was carried out at CL’s laboratory in Cambridge and contributed to interpretation of data. NV and NS contributed equally to this manuscript, carried out all the experiments, analysed the data and helped with figure preparation and drafting the manuscript. JDFL wrote the manuscript with help from all the authors.

## 7. Funding source

This work was supported by the University of Bath through a University Research Studentship to NV, the University of Bath Alumni Fund for seed funding our work on glioblastoma, and the Royal Society grant ZR-Y0113. NS is funded by a GW4 BioMed MRC Doctoral Training Partnership. We acknowledge a SMRS award from Newnham College which enable visits of NV to Cambridge to carry out live cell imaging experiments. We also acknowledge funding MRC MR/M01102X/1 to CL.

## 8. Conflict of interest

The authors declare no conflict of interest.

## 9. Acknowledgments

We are grateful to the University of Bath Imaging Facility suite for support with flow cytometry analysis and confocal microscopy. We are grateful to Dr Mark Skehel (MRC Laboratory of Molecular Biology, Biological Mass Spectrometry and Proteomics, Cambridge, UK) for mass spectrometry analysis of TRABID interactome. We thank Dr Joshua E Flack (MRC-LMB) for contributing HECTD1 KO cell lines, Professor Irene Zohn (Children’s National Research Institute, USA), for the full-length mouse Hectd1 mammalian expression vector, and Professor David J Stephens (University of Bristol) for contributing hTERT-RPE cells.

## Abbreviations

APC/c: Anaphase Promoting Complex/cyclosome
DUB: deubiquitylase
HECT: Homologous to the E6AP carboxyl terminus
NEBD: nuclear envelope breakdown
NZF: Npl14 Zinc Finger
SCF: Skp, Cullin, F-box containing complex
SAC: Spindle assembly checkpoint
UBD: Ubiquitin binding domain
UPS: Ubiquitin Proteasome System

## Notes

### Competing Interest Statement

The authors have declared no competing interest.

