## Supplementary Figures for "The E3 ubiquitin ligase HECTD1 contributes to cell cycle progression through an effect on mitosis"

Supplementary Figure 1

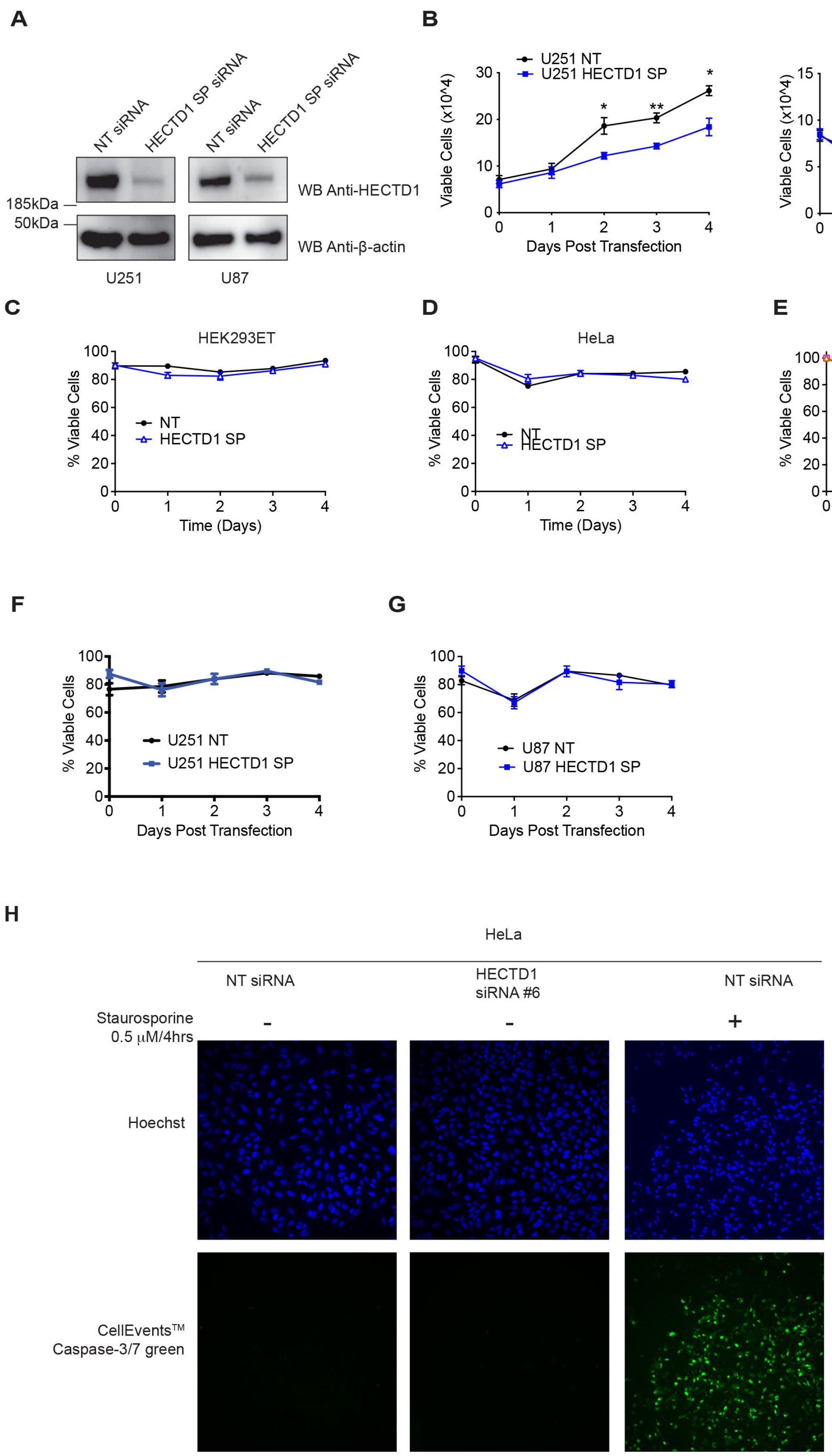


**Supplementary Figure 1. HECTD1 depletion does not trigger cell death**

**A)** Immunoblot showing HECTD1 knock down efficiency in U251 and U87 cells 48 hrs post siRNA treatment. **B)** Viable cell count (x$10^4$) is shown for U87 and U251 treated with either Non-Targeting (NT) or HECTD1 SMARTpool (SP) siRNA. Data plotted as mean with error bars that represent ±S.E.M., over three independent experiments (n=3), **p<0.01, *p<0.05 by paired student's t-test. **C-G**) Data plotted as mean of viable cells (%) with error bars that represent ±S.E.M., over three independent experiments (n=3), **p<0.01, *p<0.05 by paired student's t-test. (**C**) HEK293ET, **(D)** HeLa, (**F**) U251, (**G**) U87 treated with NT or HECTD1 SP siRNA. (**E**) Mean of viable cells for HEK293T WT, HECTD1 KO1 and KO2 plotted as above. **H**) Microscopy images obtained using an EVOS cell imaging system showing CellEvent$^{TM}$ Caspase-3/7 green detection reagent as a marker of apoptotic cell death. HeLa cells were incubated with NT or HECTD1 SP siRNA for 48 hrs, prior to detection with CellEvent$^{TM}$ Caspase-3/7 green reagent. As a positive control for this assay, HeLa cells were treated with 0.5 μM of Staurosporine for 4hrs to trigger caspase-mediated apoptotic cell death.

Supplementary Figure 2

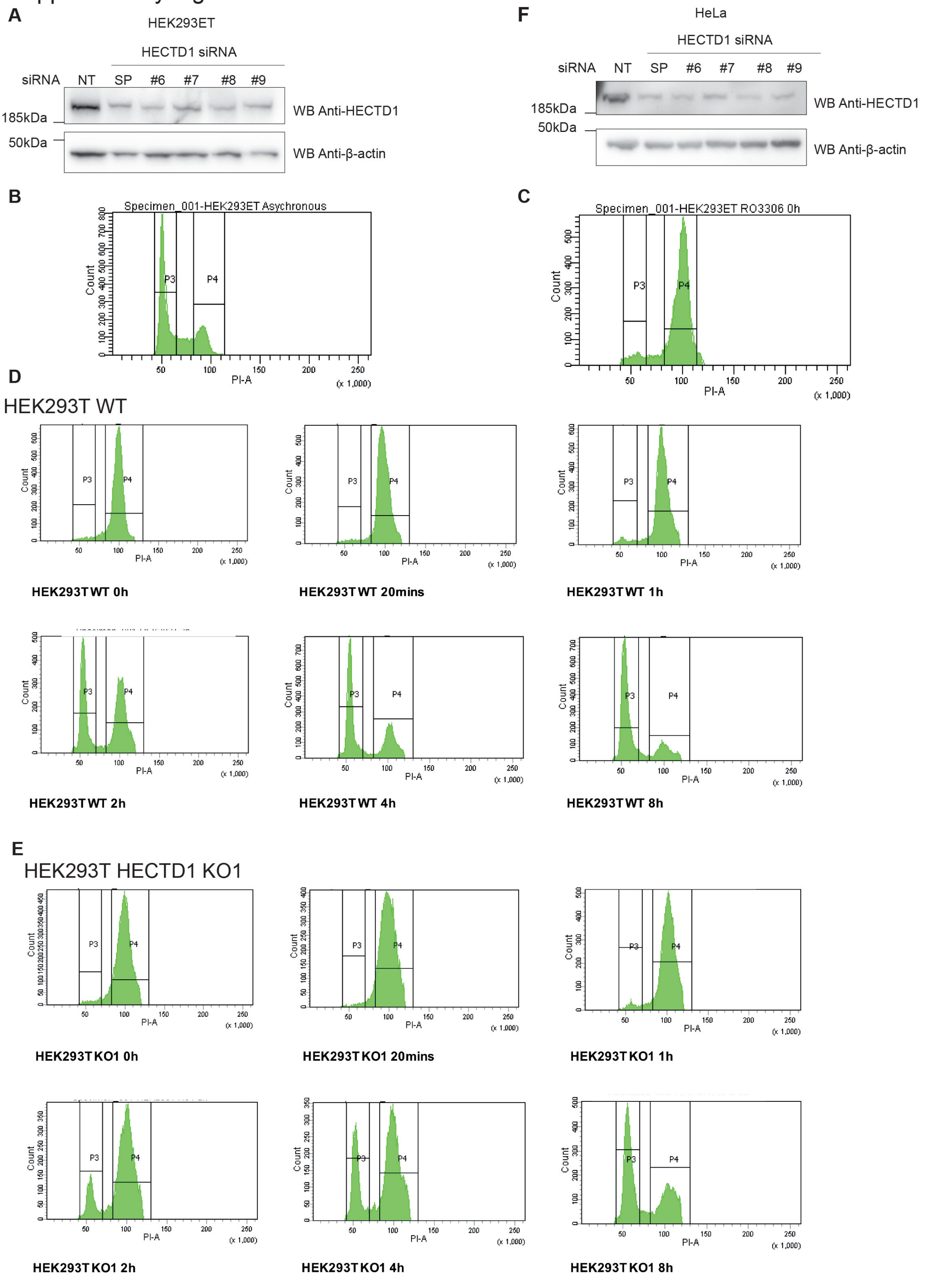


**Supplementary Figure 2. Flow cytometry cell cycle analysis of wild-type and HECTD1-depleted cells synchronised in late G2 using RO3306**

**A**) Immunoblotting showing the efficacy of HECTD1 siRNA-mediated knockdown in HEK293ET cells. Cells were transiently transfected for 48 hrs using 10 pmol, 20 pmol, or 40 pmol HECTD1

(SMARTpool, SP) siRNA with Lipofectamine 2000 per well of a 24-well plate. **B & C**) Flow cytometry cell cycle analysis of asynchronous HEK293T cells (**B**), or cells which were synchronised in late G2 with 9 µM RO3306 for 20 hrs prior to analysis (**C**). Cells were fixed using 70% ethanol, and stained using 2 µg/ml PI, with 100 µg/ml RNase A, for 30 min at room temperature. Stained samples were then analysed immediately by flow cytometry. Gated population percentages are indicated on each graph. PI-A of 50 is equivalent to 2N (G1 population), and PI-A of 100 is equivalent to 4N (G2/M population). PI-A of 50 is equivalent to 2N (G1 population), and PI-A of 100 is equivalent to 4N (G2/M population). **D, E**) Flow cytometry cell cycle analysis of HEK293T WT (**B**) and HECTD1 KO1 (**C**) following release from RO3306 block. Cells were released in full media prior to processing at the indicated time points (20 min, 1 h, 2 hr, 4 h, 8 h). **F**) Immunoblotting showing the efficacy of HECTD1 siRNA-mediated knockdown in HeLa cells using RNAiMAX as transfection reagent.

Supplementary Fig 3

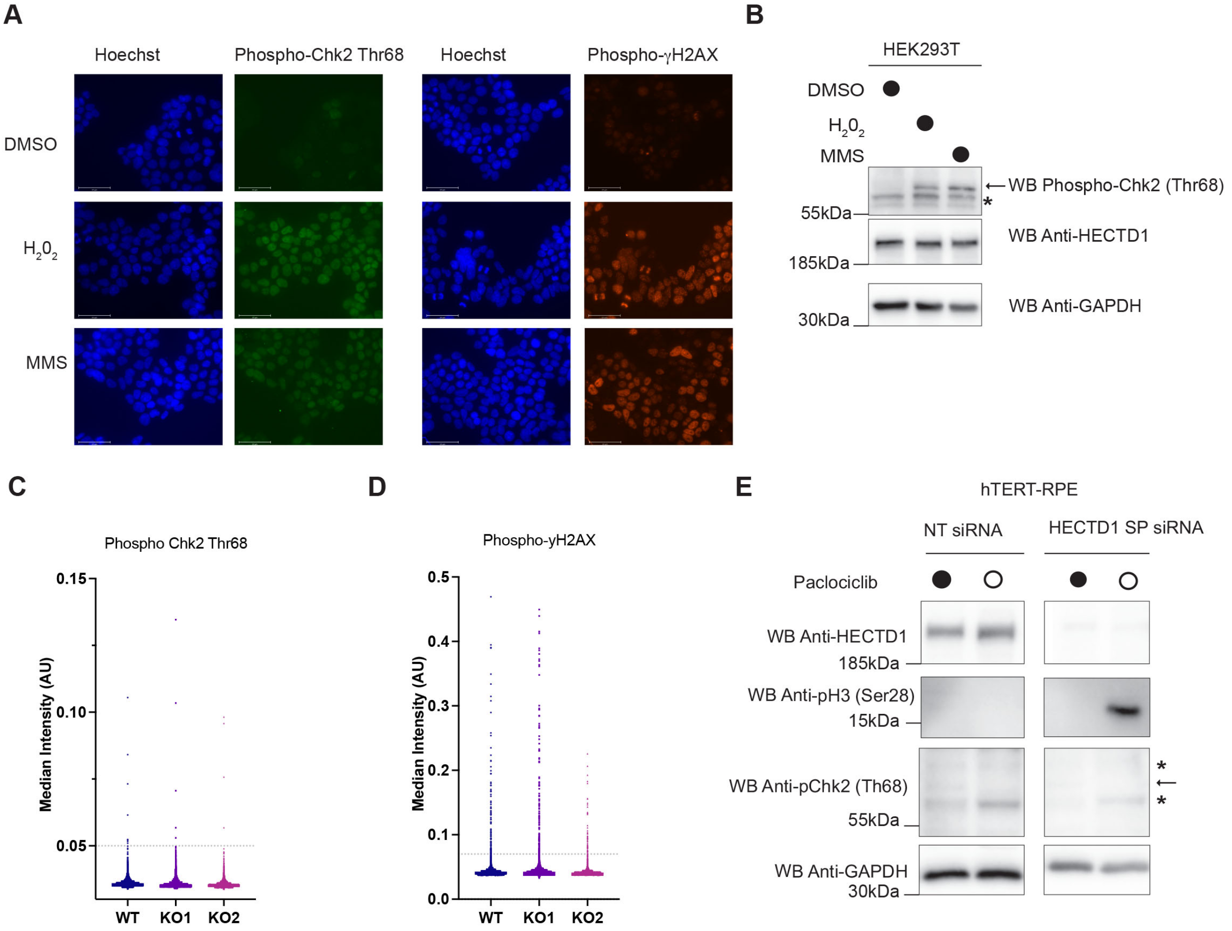


**Supplementary Figure 3. HECTD1 depletion does not induce DNA damage markers**

**A)** Microscopy images obtained using an EVOS cell imaging system validating the detection of cell with DNA damaged using phospho-Chk2 (Thr68) and phospho-γH2AX antibodies. HEK293T cells treated with 0.1 mM $H_2 0_2$ or 0.3 mg/ml of Methyl MethaneSulfonate (MMS) for 30 min prior to fixation with 4% PFA and immunostaining using phospho-Chk2 (Thr68) and phospho-γH2AX antibodies [1]. Alexa Fluor546 Donkey anti-Rabbit was used as secondary antibody and Hoechst as nuclear stain and then mounted with VectaShield Antifade Mounting Medium. **B**) HEK293T cells were treated as in A) but lysed using RIPA buffer and analysed by immunoblotting using phospho-Chk2 (Thr68). Arrow indicates phospho-Chk2 (Thr68)-specific signal while * show non-specific signals. **C-D**) High content microscopy analysis of (**C**) phospho-Chk2 (Thr68) and (**D**) phospho-γH2AX-positive cells in HEK293T wild-type, HECTD1 KO1 and KO2 cell lines. No significant difference was observed. **E**) Immunoblotting showing the increase in phospho-H3 (Ser28) levels upon siRNA mediated knockdown of HECTD1 in hTERT-RPE cells. This increase did not coincide with an increase in phospho-Chk2 (Thr68). Palbociclib (150 nM for 12 hrs) was used as control since it synchronises cells in G1 [2].

Supplementary Figure 4

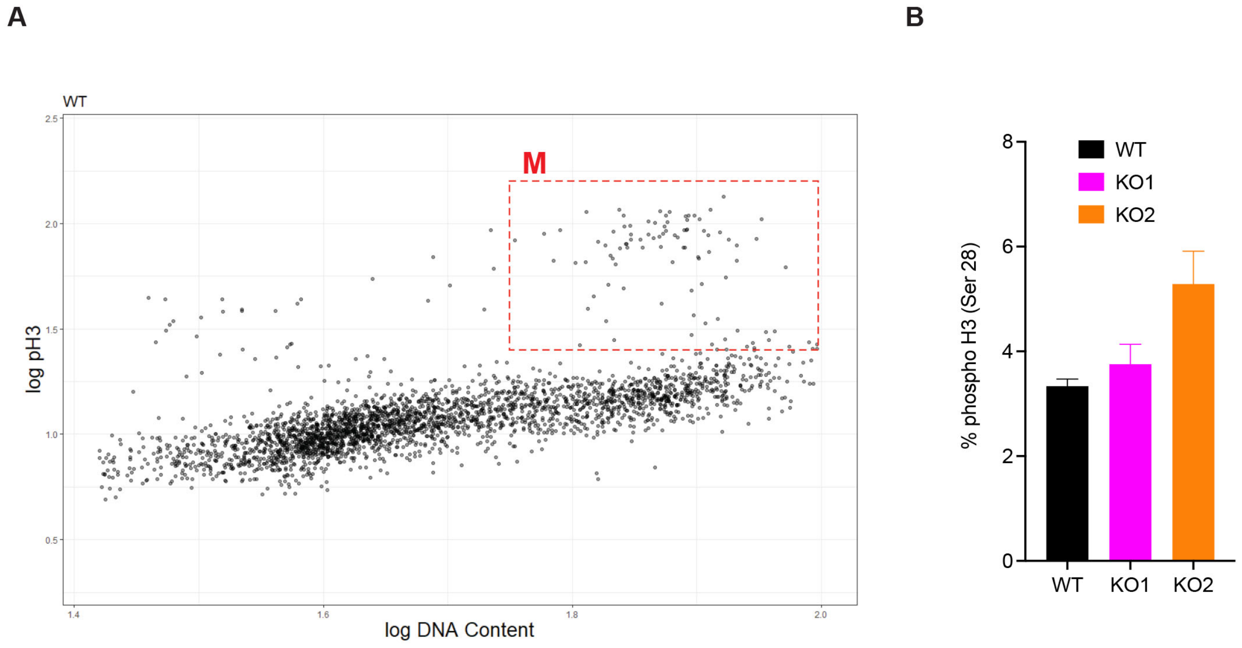


**Supplementary Figure 4. Quantification of phospho-H3 (Ser28)-positive cells**

**A)** Examples showing analysis of high content microscopy data obtained using phospho-H3 (Ser28) and Hoechst signal intensity to identify cells in M-phase. **B)** Quantification of high content microscopy data in HEK293T wild-type, KO1 and KO2 cell lines using gates set up in A). Each condition was set up as three independent wells. No significant difference was observed.

### Supplementary Figure 5

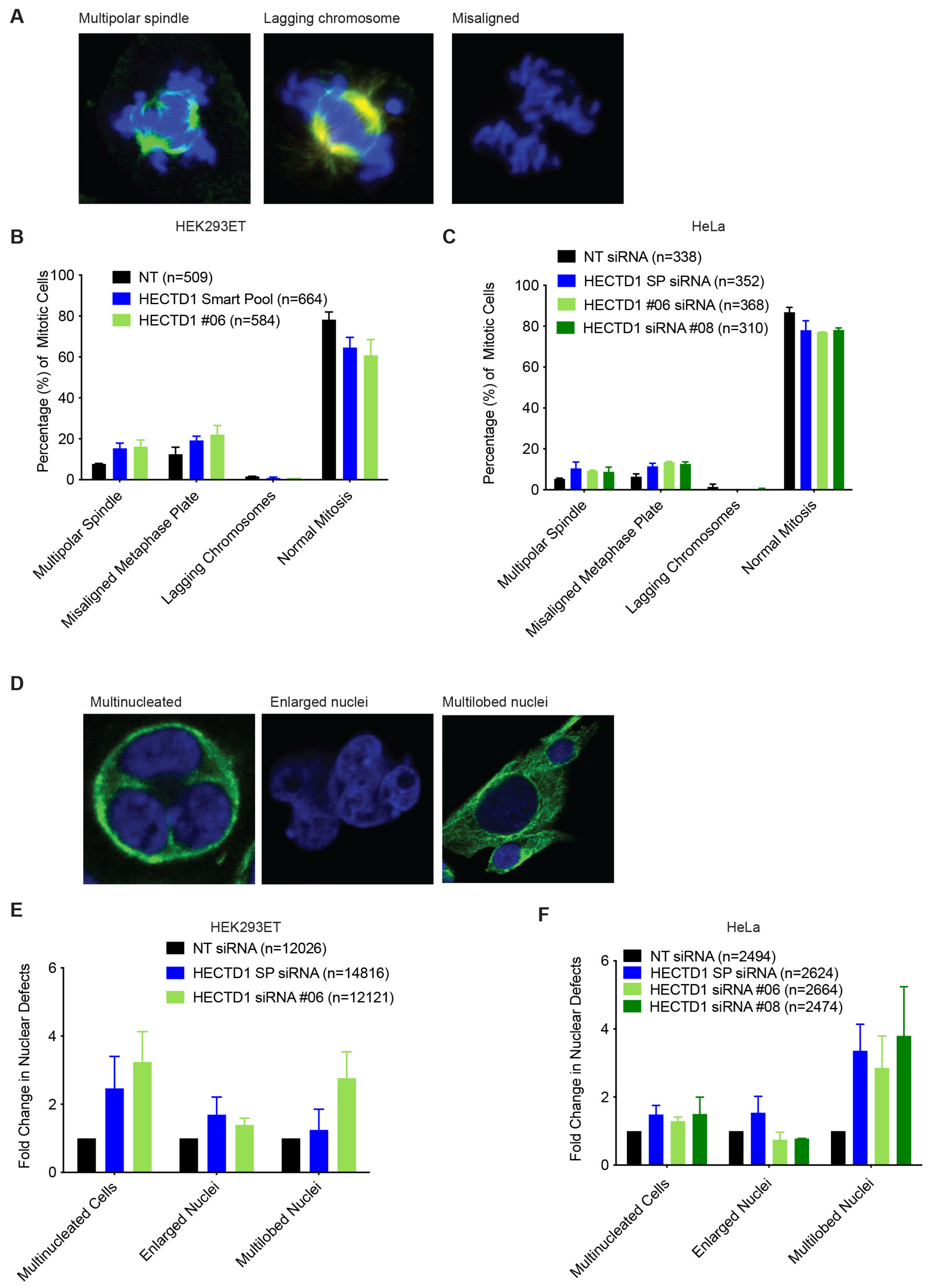


**Supplementary Figure 5. HECTD1-depleted cells show no significant mitotic defect**

A) Representative confocal images of mitotic defects scored in HeLa cells. **B**) HEK293ET and **C**) HeLa cells were scored according to chromatin morphology (Hoechst, blue) or spindle morphology based on (α-Tubulin, green) staining following 48 hrs incubation with either NT (non-targeting) siRNA, HECTD1 SMARTpool (SP) siRNA or individual HECTD1 siRNA #06, and #08. Cells were categorised into three mitotic defect phenotypes: multipolar spindle, misaligned metaphase plate, or lagging chromosomes. Cells with no observable mitotic defects were scored as cells that were in normal mitosis. Data plotted as mean with error bars that represent ±S.E.M., over 6 biological repeats. Average increase in multipolar spindle = HECTD1 siRNA of 8.0% HEK293ET; 4.1% HeLa. Average misaligned metaphase plate = HECTD1 siRNA of 7.1% HEK293ET; 6.2% HeLa. However, no statistical significance was found using a one-way ANOVA test. **D**) Representative images of each nuclear defect scored (HeLa) according to chromatin morphology Hoechst (blue). α-Tubulin is shown in green. **E**) HEK293ET and **F**) HeLa cells were scored following 48 hrs incubation with either NT (Non-Targeting) siRNA, HECTD1 SMARTpool (SP) siRNA, individual HECTD1 siRNA #06, and #08. Cells were grouped into three nuclear defect phenotypes: multinucleated cells, enlarged nuclei, or multilobed nuclei. Data plotted as mean with error bars that represent ±S.E.M., over 6 biological repeats. HEK293ET HECTD1 #08 siRNA represents one independent experiment. No statistical significance was found using a one-way ANOVA test.

### Supplementary Figure 6

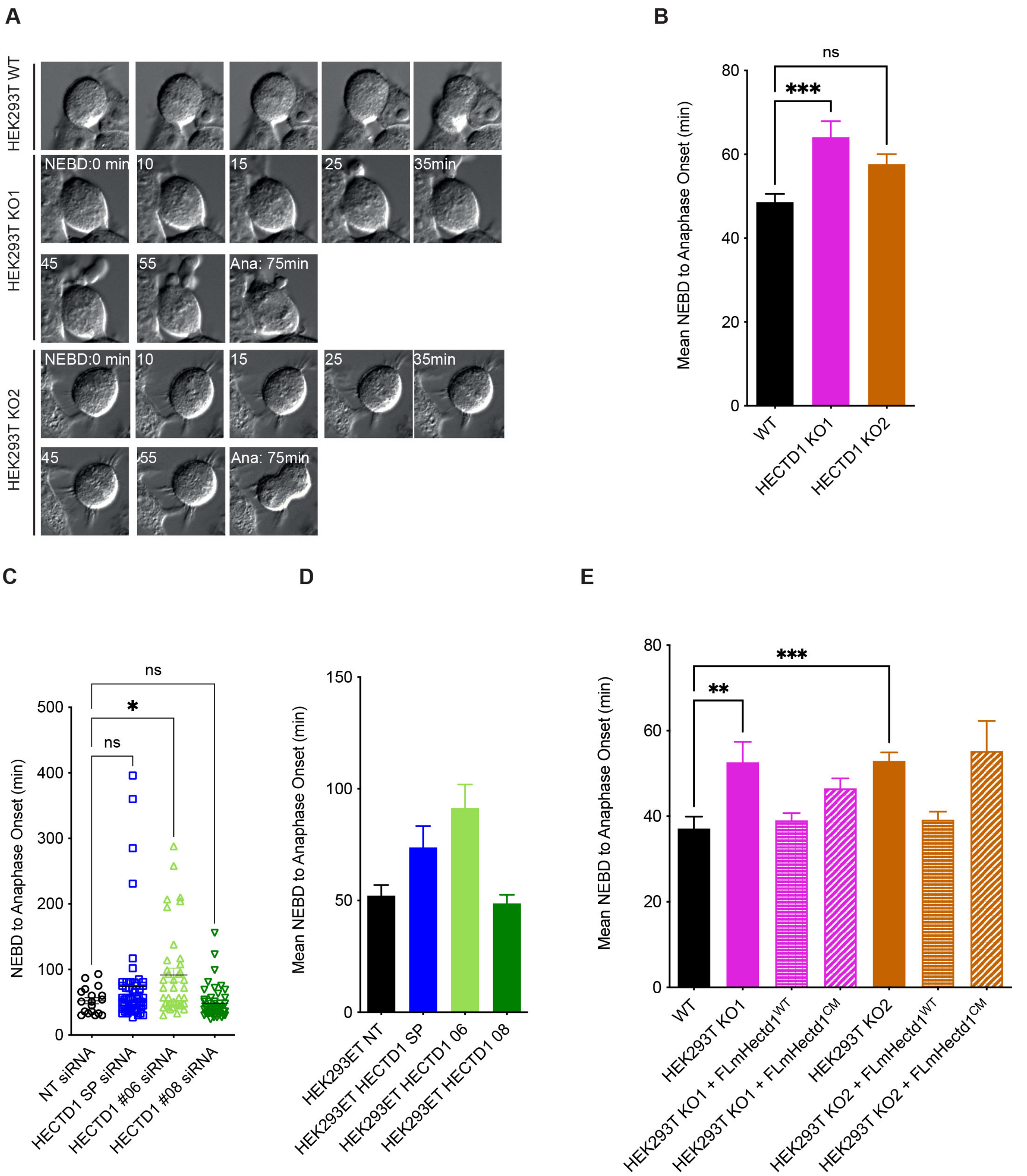


**Supplementary Figure 6. HECTD1 ubiquitin ligase activity is important for the normal timing of NEBD to anaphase onset**

**A**) Representative still images of cells scored in Fig 4D. Cells were imaged using an Olympus IX81 microscope with a 40X oil immersion objective lens and Hammatsu ORCA-ET Camera at 37°C. Micro-Manager was used to acquire and analyse images [3]. Interestingly, the anaphase image of KO1 appeared tripolar, reflecting our earlier data that some HECTD1-depleted cells might have multipolar spindles (Supplementary Fig 5B). **B**) Mean time taken (min) for HEK293T wild-type, HECTD1 KO1 and KO2 to progress from NEBD to anaphase onset for data shown in Fig 4D. Error bars represent ±S.E.M., ***$p<0.001$, using a one-way ANOVA with a Dunnett's post-test. Number of cells filmed are as follows, WT = 116, KO1 = 136, and KO2 = 161, filmed over 4 independent experiments. **C**) Mean time taken (min) for cells to progress from NEBD to anaphase onset for data shown in Fig 4D. **C**) Vertical scatter plot showing the time taken for each cell to progress from NEBD to anaphase onset 72 hrs post transfection of HEK293ET with either Non-Targeting (NT), HECTD1 SP (SMARTpool), and individual HECTD1 siRNA#06 and #08. **D**) Mean time taken (min) for cells to progress from NEBD to anaphase onset for data shown in C). Error bars represent ±S.E.M., $*p<0.05$, by using a one-way ANOVA with a Dunnett's post-test. Number of cells filmed are as follows, NT (72 hrs) = 18, HECTD1 SP (72 hrs) = 57, HECTD1 siRNA#06 (72 hrs) = 39, and HECTD1 siRNA#08 (72 hrs) = 45, filmed over 3 independent experiments. Similar data were obtained following transfection for 48 hrs (not shown). **E**) Mean time taken (min) for cells to progress from NEBD to anaphase onset for data shown in Fig 4E). Error bars represent ±S.E.M., $***p<0.001$, by a one-way ANOVA with a Dunnet's post-test. Number of cells filmed are as follows, HEK293T WT = 127, KO1 = 97, KO1 + HA-FLmHectd1$^{WT}$ = 48, KO1 + HA-FLmHectd1$^{CM}$ = 108, KO2 = 153, KO2 + HA-FLmHectd1$^{WT}$ = 46, and KO2 + HA-FLmHectd1$^{CM}$ = 19, filmed over 3 independent experiments.

Supplementary Figure 7

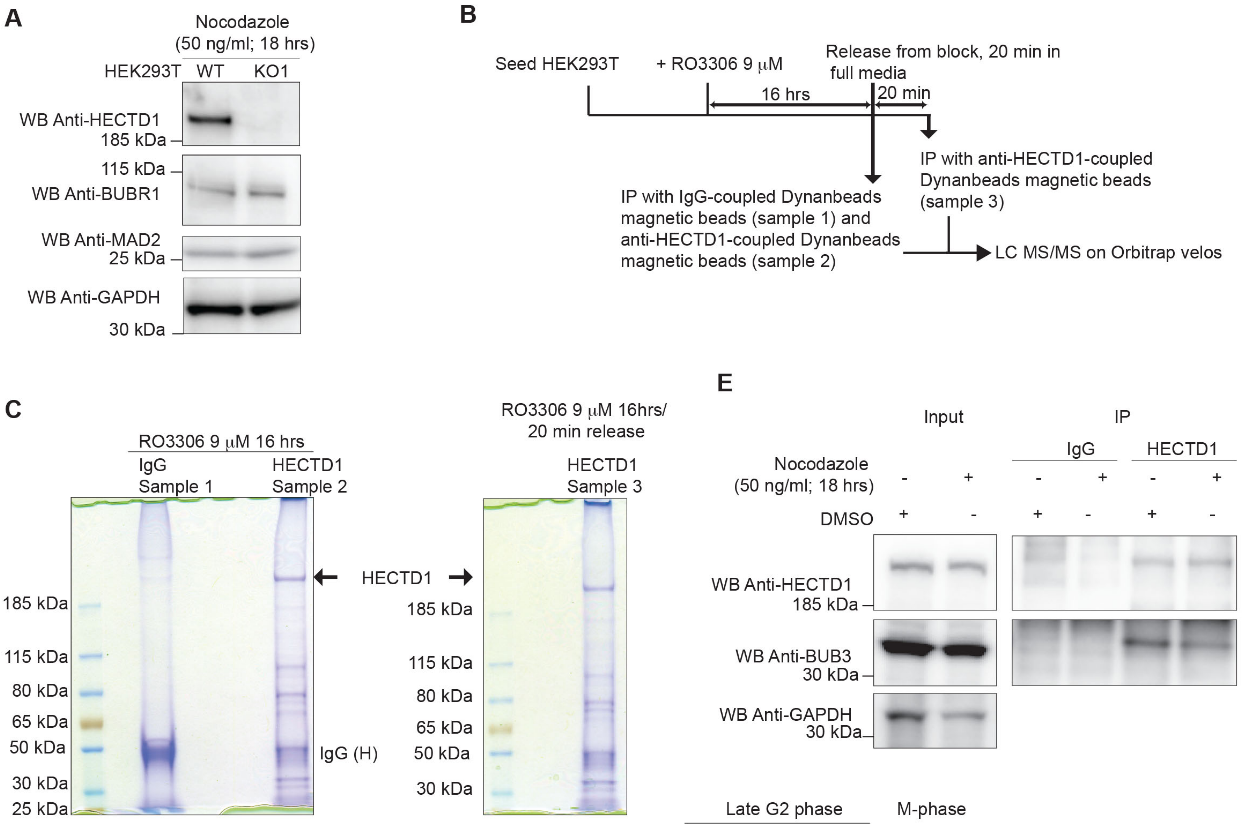


**D**

▬ Mitotic checkpoint complex (MCC)
⬢ Chromosome passenger complex (CPC)
★ Microtubule dynamics
⬬ Chromosomal assembly and segragation

| Identified Proteins | Uniprot ID | Mw (kDa) | Sample 1 (IgG IP) (Late G2 phase) | Sample 2 (HECTD1 IP) (Late G2 phase) | Sample 3 (HECTD1 IP) (M-phase) | |
|---|---|---|---|---|---|---|
| E3 ubiquitin ligase HECTD1 | HECTD1 | 289 | 0 | 82 | 68 | |
| Centromere protein F | CENPF | 368 | 0 | 42 | 2 | ★ |
| Up-reulator of cell proliferation | URGCP | 105 | 0 | 32 | 21 | |
| Centrosomal protein of 170 kDa | CEP170 | 175 | 0 | 27 | 5 | ★ |
| Cell division 5-like protein | CDC5L | 92 | 2 | 20 | 12 | |
| Structural maintenance of chromosomes protein 2 | SMC3 | 142 | 0 | 9 | 12 | ⬬ |
| Structural maintenance of chromosomes protein 3 | SMC2 | 136 | 0 | 9 | 11 | ⬬ |
| Cell growth-regulating nucleolar protein | LYAR | 44 | 0 | 6 | 10 | |
| Nucleolar and spindle-associated protein 1 | NUSAP1 | 49 | 0 | 6 | 4 | |
| **Mitotic checkpoint protein BUB3** | **BUB3** | 37 | 0 | 6 | 4 | ▬ |
| Cell division control protein 6 homolog | CDC6 | 63 | 0 | 4 | 0 | |
| Centromere protein V | CENPV | 30 | 0 | 4 | 2 | ★ |
| Condensin-2 omplex subunit G2 | NCAPG2 | 131 | 0 | 3 | 4 | |
| Borealin | Borea | 31 | 0 | 2 | 5 | ⬢ |
| Cyclin-dependent kinase 13 | CDK13 | 165 | 0 | 3 | 2 | |
| p21-activated protein kinase-interacting protein 1 | PAK1IP1 | 44 | 0 | 2 | 3 | |
| Inner centromere protein | INCENP | 105 | 0 | 2 | 0 | ⬢ |
| Cell division cycle-associated protein 2 | CDCA2 | 113 | 0 | 0 | 2 | |
| Centromere protein U | MLF1P | 48 | 0 | 2 | 0 | ★ |
| Nuclear mitotic apparatus protein 1 | NUMA1 | 238 | 0 | 0 | 25 | ★ |
| Condensin-2 complex subunit D3 | NCAPD3 | 169 | 0 | 0 | 8 | |
| S phase cyclin A-associated protein in the ER | SCAPER | 158 | 0 | 0 | 3 | |
| Sister chromatid cohesion protein PDS5 homolog B | PDS5B | 165 | 0 | 0 | 5 | |
| Cyclin-dependent kinase 1 | CDK1 | 34 | 0 | 0 | 2 | |
| Cyclin-dependent kinase 9 | CDK9 | 43 | 0 | 0 | 2 | |
| Centromere protein h, isoform CRA_b | CENPH | 26 | 0 | 0 | 3 | |
| Major centromere autoantigen B | CEPNB | 65 | 0 | 0 | 2 | ★ |
| Centromere protein Q | CENPQ | 31 | 0 | 0 | 2 | ★ |
| Centromere protein P | CENPP | 33 | 0 | 0 | 2 | ★ |
| Cyclin-dependent kinase 12 | CDK12 | 164 | 0 | 0 | 2 | |

Number of unique peptides

**Supplementary Figure 7. Proof of principle study of HECTD1 interactome in late G2 and M-phases**

**A)** Immunoblot analysis of wild-type and HECTD1 KO1 HEK293T cells treated for 18 hrs with 50 ng/ml to activate the SAC. While wild-type cells remain in mitosis upon SAC activation as shown by the pH3 (Ser28) signal, HECTD1-depleted cells show a decrease in signal intensity suggesting that these cells might not be efficiently blocked in mitosis. **B**) Schematic of the proteomics experiments to identify candidate interactors of endogenous HECTD1 in HEK293T. **C**) Coomassie stained gel showing proteins captured using IgG-coupled magnetic beads (G2), anti-HECTD1-coupled magnetic beads, from lysates of cells synchronised in late G2 and M-phase (See Supplementary Figure 2B & C for validation of synchronisation). Each lane was cut in 24 gel slabs which were analysed by LCMS/MS using an Orbitrap velos (n = 1 IP performed for each sample). **C**) Selected curated list of cell cycle related HECTD1 candidate interactors. **D**) Immunoprecipitation assay showing endogenous interaction between HECTD1 and BUB3 HEK293T cells. This interaction is not dependent upon SAC activation as it is also detected in the DMSO-treated sample. SAC activation was carried out as in Fig 6A. Input samples are shown and GAPDH is used as loading control. **E)** immunoprecipitation assay showing that the HECTD1-BUB3 interaction is not induced by Nocodazole treatment suggesting it is independent of SAC activation.

Supplementary Figure 8

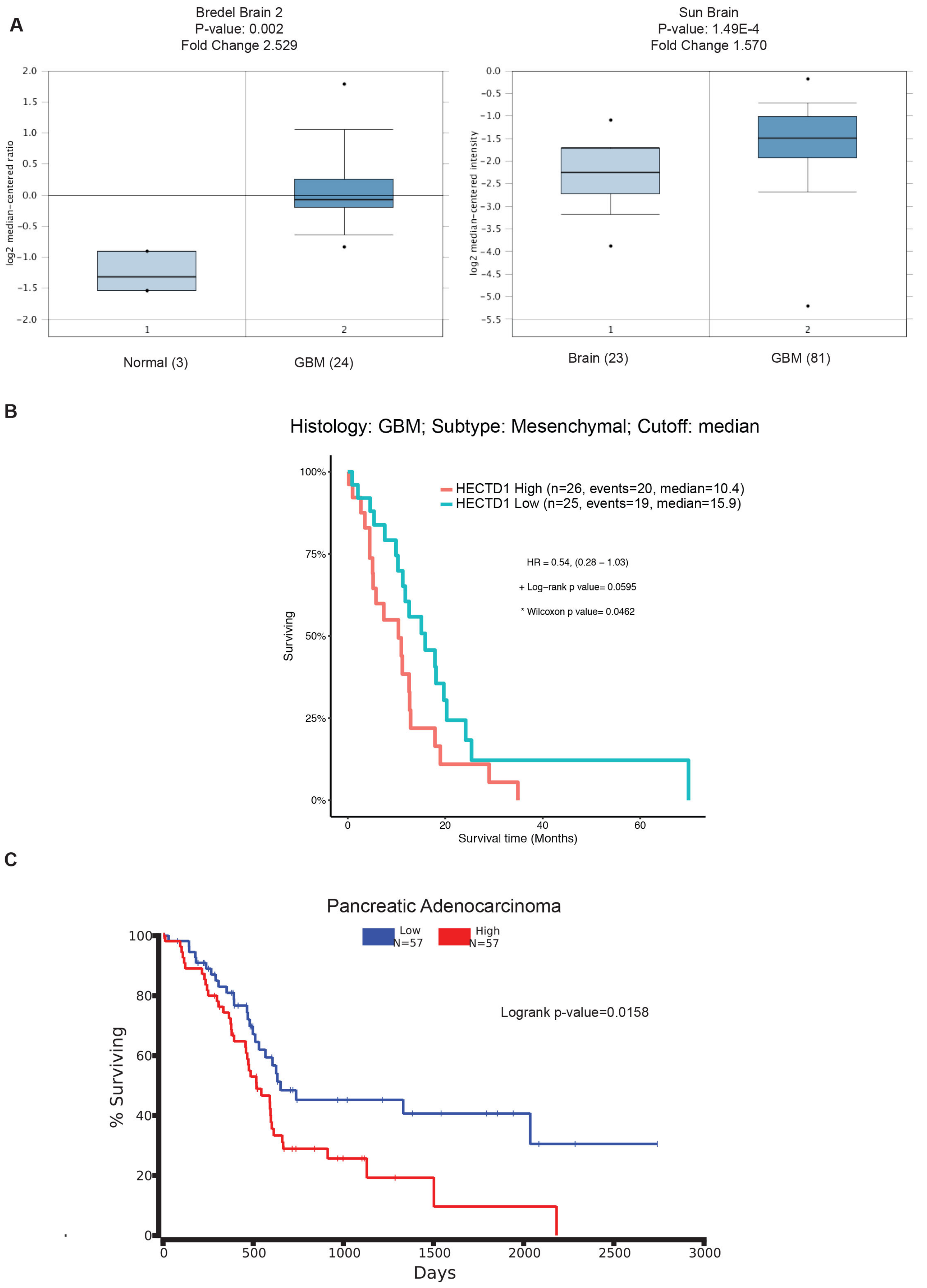


**Supplementary Figure 8. HECTD1 and cancer**

**A**) ONCOMINE[TM] ([https://www.oncomine.org/resource/login.html)](https://www.oncomine.org/resource/login.html) showing the fold change in *HECTD1* mRNA levels in normal vs Glioblastoma samples, in the Bredel Brain 2 and the Sun Brain datasets. **B-C**) TCGA analysis shows high HECTD1 mRNA expression correlates with lower survival time in GBM (Mesenchymal subtype) (**B**) and in pancreatic adenocarcinoma **C**).
